# spSheaf: sheaf-theoretic spatial domain inference from raw molecules without cell segmentation

**DOI:** 10.64898/2026.11.09.765244

**Authors:** Eduardo Paluzo-Hidalgo, Kazumitsu Maehara, Yusuke Imoto

**Author notes:** Contributing authors.

## Abstract

Spatial-transcriptomics pipelines usually require cell segmentation and discard 20–60% of transcripts. We introduce spSheaf, a segmentation-free spatial domain inference method built on cellular sheaf theory. Within a domain, neighbouring regions have compatible local expression structure; across a transition between programs they remain discordant after alignment of their local subspaces. spSheaf quantifies this alignment-adjusted disagreement and attenuates transitions during clustering. On the MERFISH hypothalamus benchmark it attains mean ARI **0.571** and NMI **0.655**, ranks first of twelve on NMI, and leads SSAM, Points2Regions and FICTURE by **0.16** ARI on held-out slices. On three further annotated sections, every method tuned, it leads on one, ties on one and trails SSAM by **0.06** ARI. The sheaf weighting adds **+0.102** ARI where domains are distinct programs and is neutral on laminar cortex. One domain above the annotated count, it divides the BST into the territories of two atlas neuronal populations on both adjacent sections.

## 1 Introduction

Spatial transcriptomics technologies now measure gene expression at submicron resolution over large tissue areas [1–3]. Imaging-based platforms such as MERFISH [4], seqFISH+ [5], osmFISH [6] and Xenium [7] detect individual RNA molecules with known coordinates and gene identities; sequencing-based technologies such as Seq-Scope [8], Stereo-seq [9] and Pixel-seq [10] reach comparable resolution through spatially resolved barcoding.

A central analytical task is *spatial domain inference*: partitioning the tissue into regions with coherent gene expression programs. Conventional pipelines first segment the tissue into cells, then cluster a cell-by-gene matrix [11, 12]. Segmentation requires auxiliary imaging, depends on morphological assumptions that fail for irregular or densely packed cells, and discards the transcripts outside segmented boundaries, often 20–60% of the total [13]. A benchmark of eleven segmentation methods on fifteen datasets from five platforms finds no method that performs best across platforms, tissues and criteria, with transcripts misassigned between neighbouring cells of different types in up to 10–15% of cells and the highest contamination on MERFISH, and concludes that segmentation sets the upper bound on every downstream analysis [14]; for a task defined at the scale of tissue regions, this is a source of method- and platform-dependent error that the task does not require. Several methods bypass it. FICTURE [13] infers pixel-level factors with a spatially coupled latent Dirichlet allocation model; SSAM [15] estimates an mRNA density field by kernel smoothing and derives cell types and domains from it; Points2Regions [16] clusters molecular point data directly; Sainsc [17] reimplements SSAM’s kernel-density cell-type mapping for in situ capture data and stops at the cell-type map. The first three are the closest comparators to the method introduced here and we benchmark against all of them under a matched protocol. Graph neural networks (GraphST [18], STA-GATE [19]), Bayesian models (BayesSpace [20], Baysor [21]), spatial autocorrelation statistics (SpaGCN [22]) and persistent homology [23] provide complementary strategies. A benchmark of 13 spatial clustering methods [24] found that none consistently dominates and that single-molecule imaging data, where the input is raw molecule coordinates rather than a matrix, remain challenging; two subsequent benchmarks reach the same conclusion [25, 26].

Here we introduce **spSheaf**. It is based on the idea that neighbouring regions within the same spatial domain exhibit compatible local expression structure, whereas transitions between expression programs remain discordant even after their local low-dimensional expression subspaces are aligned. The construction consists of a local principal-component basis at every point of a spatial grid, a singular-value-decomposition alignment of adjacent bases, and the residual between the two points’ transported coordinates after alignment; in the language of cellular sheaves [27], these are the stalks, the restriction maps and the edge disagreement. spSheaf quantifies this alignment-adjusted local disagreement and uses it to reduce the influence of transitional or heterogeneous regions during clustering, producing spatial domains directly from raw molecule data. Unlike spectral methods based on the sheaf [27] or connection [28] Laplacian, which smooth the boundary signal through eigendecomposition, it targets the *discontinuities* of the sheaf structure; the construction shares its local bases and alignment with vector diffusion maps [28] and ddHodge [29] and differs in base space, coordinates and downstream operation (Discussion).

We validate spSheaf on the Moffitt et al. (2018) MERFISH hypothalamus benchmark [30], where it ranks first on NMI among twelve methods and leads SSAM, Points2Regions and FICTURE on identical molecule tables under each of the three model-selection rules the five slices allow; on three further annotated cortical sections — osmFISH [6] and two SpaceTx [31] sections not used in development, an Allen Institute prototype MERFISH and a BaristaSeq [32] section — with every method tuned on its own grid; and, through a feature-fiber variant, on three-dimensional subnuclear data [33] and multiplexed protein imaging [34], as demonstrations of applicability for which no accuracy claim is made.

## 2 Results

### 2.1 Overview of the spSheaf method

spSheaf takes a set of molecule detections 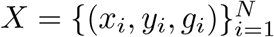, coordinates and gene identity, and returns a partition of the tissue into *k* domains without cell segmentation or auxiliary imaging (Fig. 1). The pipeline has eight steps: (1) *expression fibers* — a grid of spacing *δ* over the tissue, at each point a gene-expression vector by Gaussian kernel smoothing of nearby molecules, normalised and log-transformed; (2) a symmetric *k*-nearest-neighbour *spatial graph* on the grid; (3) *stalks* — at each vertex a local PCA basis of the locally centred fibers of its neighbourhood, the vertex’s coordinates being its own fiber’s scores in that basis; (4) *restriction maps* — for each edge, the singular value decomposition of the stalk overlap, which carries both vertices’ coordinates into one principal-angle-aligned frame; (5) *sheaf disagreement* — the squared difference of the transported coordinates; (6) *boundary score* — the per-vertex mean disagreement over incident edges, Gaussian-smoothed; (7) *boundary-aware clustering* — expression features attenuated by an exponential weight in that score, concatenated with scaled coordinates, clustered by *k*-means; (8) *cleanup* of small fragments (Methods). No sheaf Laplacian is assembled or diagonalised.

**Fig. 1.**
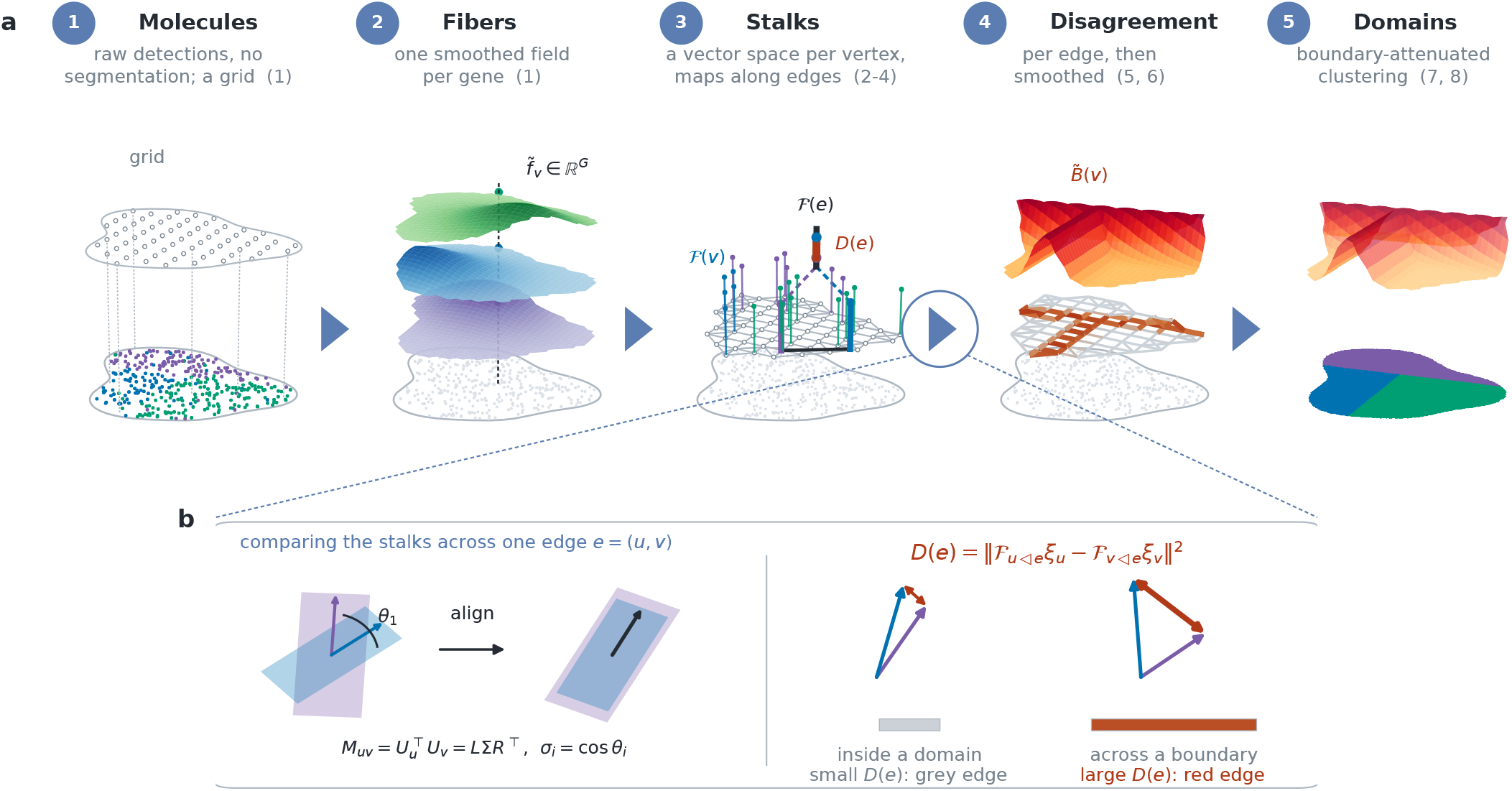
spSheaf, from molecules to domains. Schematic; nothing here plots measured data. **a**, The pipeline as one sequence, left to right, each stage on the same tissue plane with its construction stacked above. *1*, molecule detections, used as they are: no cell segmentation. *2*, Gaussian smoothing turns each gene into a continuous field over the plane, one per gene; reading up the vertical at a grid point *v* gives that vertex’s expression fiber 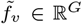 (step 1). *3*, a *k*-nearest-neighbour graph over the grid carries a cellular sheaf: over each vertex a stalk ℱ (*v*), the *d* leading principal directions of its neighbours’ fibers, drawn as a fiber standing on the base, with restriction maps along the edges (steps 2–4). *4*, each edge carries its disagreement *D*(*e*) — grey where the two stalks agree, red where they do not — and the per-vertex mean over incident edges is Gaussian-smoothed into the field 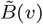 above (steps 5, 6). *5*, expression features attenuated by that field, concatenated with scaled coordinates and clustered, give the partition; the ridges of the field lie over its borders (steps 7, 8). Numbers in parentheses are the eight steps of Section 2.1. **b**, A magnification of the step ringed in **a**, which carries stage 3 into stage 4: how one edge gets its score. The overlap 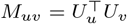 factors as *L*σ*R*^*⊤*^, whose singular values are the cosines of the principal angles between the two stalks; the restriction maps ℱ_*u⊴e*_ = *L*^*⊤*^ and ℱ_*v⊴e*_ = *R*^*⊤*^ carry both into one frame, and *D*(*e*) is the residual between the transported coordinates *after* that alignment. Within a domain they nearly coincide; across a boundary they do not.

### 2.2 Benchmarking on the MERFISH mouse hypothalamus

The Moffitt et al. (2018) MERFISH dataset [30] is a standard benchmark [24]: 135 genes, five annotated slices of the preoptic region (Bregma 0.04–0.24; 5,488–5,926 annotated cells each, eight anatomical domains), 1.8–2.0 million molecules per section, all of which every method reads (Section 4.1). spSheaf reaches mean ARI 0.571 and NMI 0.655 over the five slices at one configuration (per-slice ARI 0.478–0.656; Supplementary Table 1, Supplementary Fig. 1). Rebuilding the comparison of Yuan et al. [24] from its source data, spSheaf ranks **first of twelve on NMI** (pooled median 0.651; next best 0.565), first on homogeneity and CHAOS, second on completeness, and fourth and fifth on silhouette width and PAS (Extended Data Fig. 1a,b). ARI is not among that figure’s released metrics; the marker-gene autocorrelation metrics are omitted because, recomputed from molecules, the annotation itself scores below every baseline on them (Moran’s I 0.109 against 0.153–0.245), their absolute values depending on cell-level preprocessing that cannot be reproduced from molecules. The only stochastic step is the *k*-means initialisation, and at *k* = 8 its effect is negligible: over 20 seeds and five slices ARI is 0.571 ± 0.000 (per-slice s.d. ≤0.001; Extended Data Fig. 1c).

**Table 1.**
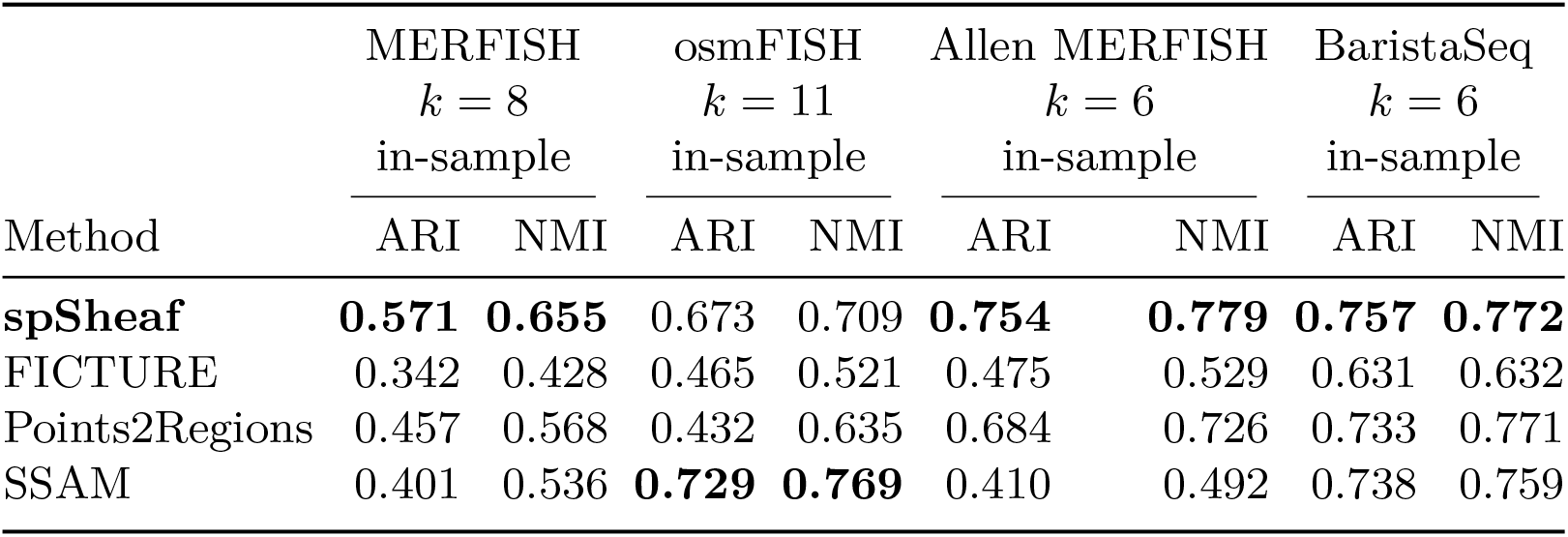
Comparison with segmentation-free methods on the same sections, every method tuned. All four are distributed with settings their authors selected, so each is given its own parameter search and selected on the annotation of the data it is scored on: SSAM over 72 configurations per MERFISH slice, 144 on osmFISH and 287–288 on the SpaceTx sections; FICTURE over 18; Points2Regions over 45 (MERFISH), 32 (osmFISH) and 24 (SpaceTx); spSheaf over 54 on MERFISH, 240 on osmFISH (108 per coordinate reference plus the *d* extension of Section 4.11) and 162 on the SpaceTx sections (Section 4.11). One rule defines a configuration for every method: a single run at a fixed seed, then the argmax over configurations. On MERFISH the configuration is the one with the highest mean ARI over the five slices, applied to all five, and the value is its mean (spSheaf: the 20-seed mean of Supplementary Table 1, seed spread *≤* 0.001); the per-slice argmax and leave-one-slice-out rules are in Fig. 2c. All four methods read the identical molecule table of each section (Section 4.1). The coordinate reference of Section 4.5 is one of spSheaf’s tuned parameters; its default is the tissue reference, which is the selected value on the three single sections, and the local reference is selected on MERFISH (Supplementary Table 3). Bold marks the best value in each column. An ARI and the NMI beside it are always the same run. On BaristaSeq the three leading ARI values lie within Points2Regions’ own seed spread and are not a separation.

|  | MERFISH |  | osmFISH |  | Allen MERFISH |  | BaristaSeq |  |
|  | *k* = 8 |  | *k* = 11 |  | *k* = 6 |  | *k* = 6 |  |
|  | in-sample |  | in-sample |  | in-sample |  | in-sample |  |
| Method | ARI | NMI | ARI | NMI | ARI | NMI | ARI | NMI |
|---|---|---|---|---|---|---|---|---|
| **spSheaf** | **0.571** | **0.655** | 0.673 | 0.709 | **0.754** | **0.779** | **0.757** | **0.772** |
| FICTURE | 0.342 | 0.428 | 0.465 | 0.521 | 0.475 | 0.529 | 0.631 | 0.632 |
| Points2Regions | 0.457 | 0.568 | 0.432 | 0.635 | 0.684 | 0.726 | 0.733 | 0.771 |
| SSAM | 0.401 | 0.536 | **0.729** | **0.769** | 0.410 | 0.492 | 0.738 | 0.759 |

### 2.3 Direct comparison with segmentation-free methods

The methods closest to spSheaf in input and intent are SSAM [15], Points2Regions [16] and FICTURE [13]. Each was run on the identical raw molecule table, its native output — a cell-type map, a molecule-level partition, a pixel-level factor map — mapped to a label per annotated cell, and all four scored against the same annotation with the same metric implementations (Section 4.11). All four are distributed with settings their authors selected, so each, spSheaf included, is given its own parameter grid and selected on the annotation, under one definition of a configuration: a single run at a fixed seed (Section 4.12).

With five slices, three selection rules can be applied identically to all four, and they differ by more than the methods do (Fig. 2c). Under the permissive per-slice argmax, the rule the competitors’ published values come from, spSheaf reaches mean ARI 0.584, Points2Regions 0.510, SSAM 0.464 and FICTURE 0.353; under one configuration chosen by mean ARI and applied to all five, the rule of Table 1, 0.571, 0.457, 0.401 and 0.342; under leave-one-slice-out, where no method sees the slice it is scored on, 0.552, 0.388, 0.343 and 0.342. The ranking is the same under all three, with margins of 0.074, 0.114 and 0.163 ARI over the best competitor. spSheaf’s grid on MERFISH is 54 configurations, its three tuned parameters at the local and at the tissue coordinate reference (Section 4.14); every rule selects the local reference here, and the one configuration selected is the one fixed during development. Per-slice tuning adds 0.013 to it, against 0.053 for Points2Regions and 0.063 for SSAM. Held out, spSheaf leads on four slices and trails Points2Regions on the fifth by 0.007 (Fig. 2a,b). SSAM’s published recipe, as implemented in SSAM 1.1.3, returns 5–7 cell types per section and only ARI 0.131 held out; the values above sweep its clustering resolution as a hyperparameter, which returns 39–63 types, the order the SSAM paper reports (Section 4.11). At the method’s default, the tissue reference, the three rules give 0.564, 0.548 and 0.540, ahead of every competitor under each.

**Fig. 2.**
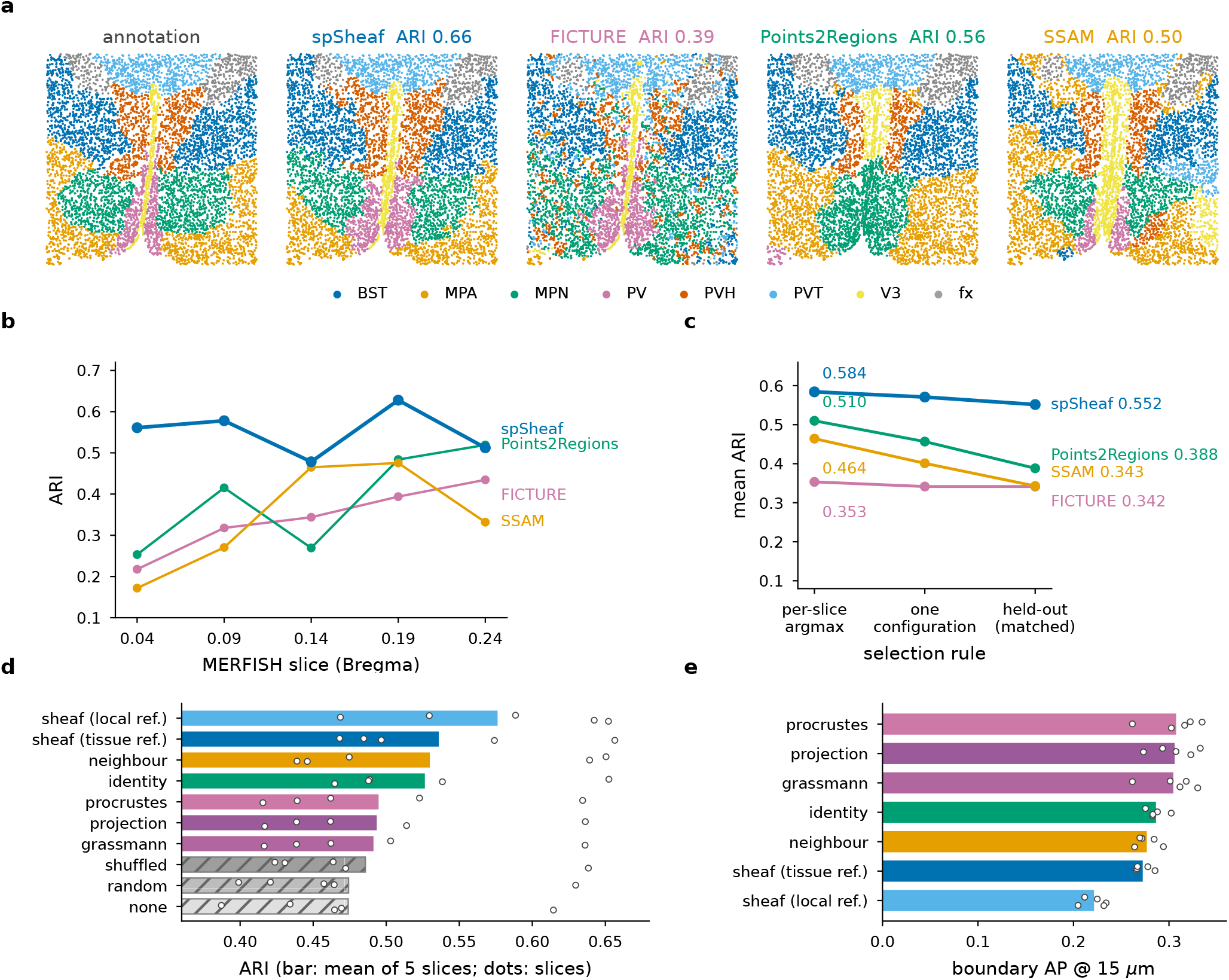
Comparison with segmentation-free methods, and the contribution of the sheaf term. **a**, Slice 0.19 of the MERFISH benchmark, per annotated cell: the annotation and the domains returned by each method, with labels Hungarian-matched to the annotation. Each competitor is drawn at its best configuration *on this slice* (the per-slice argmax of panel **c**, i.e. the most favourable setting for that method); spSheaf is drawn at its one configuration of Table 1. The ARI of the drawn run is given above each map. **b**, Per-slice ARI on the five MERFISH slices, every method’s hyperparameters selected by leave-one-slice-out from its own grid (spSheaf: the 54-configuration grid of Section 4.14); all are single runs at the held-out-selected configuration. **c**, Mean ARI under the three selection rules the five slices allow, applied identically to all four methods: *per-slice argmax*, a different configuration per slice chosen on that slice’s own annotation; *one configuration*, chosen by mean ARI over the five and applied to all five, the rule of Table 1; and *held-out*, the rule of panel **b. d**, Edge-score ablation: ten arms differing only in the edge score *D*(*e*); *sheaf (local ref*.) is the method, *sheaf (tissue ref*.) the same score with the optional tissue-referenced coordinates. Bars are means over 5 slices *×*5 *k*-means seeds, scored at grid level (which is why the method arm reads 0.576 rather than the cell-level 0.571 of Table 1); open circles are the five per-slice means. Hatched bars are the three null controls; *none* is the unweighted pipeline (*α*_*b*_ = 0). **e**, The same fields evaluated against annotated boundaries, as average precision at a 15 *µ*m tolerance; bars are five-slice means, circles the per-slice values.

On the three single sections there is no slice to hold out, so Table 1 reports every method at the argmax of its own grid, selected on the annotation it is scored against — an upper bound for each. spSheaf’s grid includes the coordinate reference of Section 4.5; the argmax is the tissue reference, the method’s default, on all three single sections, and the local reference on MERFISH. spSheaf leads on the Allen prototype (0.754 against 0.684) and BaristaSeq (0.757; the three leading values lie within Points2Regions’ seed spread and are a tie) and trails SSAM on osmFISH by 0.056 (0.673 against 0.729; Sections 2.6, 2.5). Points2Regions varies with its seed (s.d. 0.026– 0.077 per MERFISH slice, 0.045 on osmFISH, against 0.001 for spSheaf); FICTURE scores higher with 20 factors aggregated into domains than with its factor count set to *k*, and the better regime is reported. Without tuning, with every parameter fixed on the MERFISH benchmark and carried over, spSheaf is 0.025 (BaristaSeq), 0.147 (osm-FISH) and 0.143 ARI (Allen prototype) behind each section’s best tuned competitor (Extended Data Table 1).

### 2.4 Contribution of the sheaf term

To separate the contribution of the sheaf score from that of weighting in general, the edge score was replaced by nine alternatives with the rest of the pipeline fixed, at the configuration selected on this benchmark (local reference), on five slices and five seeds (Fig. 2d; Section 4.9). Three arms are null controls: the field permuted across vertices, an independent random field, and no weighting. The sheaf score is first on ARI and NMI, by +0.102 ARI over no weighting, +0.102 over a random field and +0.090 over its own spatial shuffle, which keeps the field’s distribution and randomises its arrangement. Against the kernel baseline that compares raw neighbouring fibers with no sheaf structure the margin is +0.046. The two components of the score contribute as follows: with tissue-referenced coordinates the principal-angle alignment is worth +0.010 over identity restrictions (0.536 against 0.526), and referencing the coordinates to the stalk’s own neighbourhood rather than the tissue mean (Section 4.5) a further +0.040 (0.576). The three scores built from subspace geometry alone — Grassmann distance, projection distance, Procrustes residual — score 0.491–0.495, against 0.474 for the random field. The alignment-adjusted disagreement of local *coordinates* accounts for the improvement; the angle between subspaces alone does not.

Evaluated directly against the annotated boundaries (average precision at 15 *µ*m; Section 4.9), the ordering reverses: the three subspace scores lead (AP 0.305–0.308, localisation error 57 *µ*m) and the method’s own score is last (0.221, 160 *µ*m; Fig. 2e). The two rankings track the width of each field’s response: the subspace scores rise over a narrow band at the interface, the coordinate scores over the full transition between programs, which is the extent the weight should attenuate. Boundary localisation and partition quality therefore order the candidate scores differently (Discussion). The same ten arms on the two SpaceTx sections at their selected configurations, where the tissue reference is the default in use, give a different picture (Supplementary Fig. 3): on both laminar sections every edge score, the method’s included, lies within seed noise of the kernel baseline (0.719 against 0.721 on the Allen prototype; 0.734 against 0.736 on BaristaSeq, where the random field and the shuffle give 0.734 and 0.737 and the Procrustes residual 0.748), the weighting itself adds 0.012–0.024 ARI, and the local-reference score, which the tissue reference replaces on these sections, falls to 0.458 on the Allen prototype. The contribution of the alignment-adjusted score is therefore conditional on the tissue: on the hypothalamus, whose nuclei are distinct expression programs with sharp interfaces, it is +0.046 ARI over kernel smoothing; on cortex organised as a compositional gradient it matches kernel smoothing (Discussion). On the benchmark the unweighted grid pipeline alone scores 0.474, already above the three molecule-level comparators under the matched rule of Table 1.

### 2.5 osmFISH mouse somatosensory cortex

The osmFISH somatosensory cortex [6] has fewer genes and more domains than MER-FISH: 33 genes, 4,839 annotated cells, 11 regions. At the argmax of its grid (tissue reference; Table 1) spSheaf reaches ARI 0.673 and NMI 0.709, and 0.658 / 0.706 at the argmax of the local-reference half (*h* = 32.5 *µ*m, *σ*_*s*_ = 19.5 *µ*m, *d* = 9; 0.648 ± 0.012 over 20 initialisations; Fig. 3), scored in a molecule frame registered without the labels (Section 4.10). Nine of the eleven regions are recovered at recall 0.72–1.00; Layer 3–4, a 73 *µ*m-wide transitional zone of 131 cells, and Layer 2–3 lateral, which differs from Layer 2–3 medial by position more than by expression, are missed at every configuration of the grid and in every initialisation. Naive *k*-means reaches ARI 0.19–0.22 on coordinates, expression or both.

**Fig. 3.**
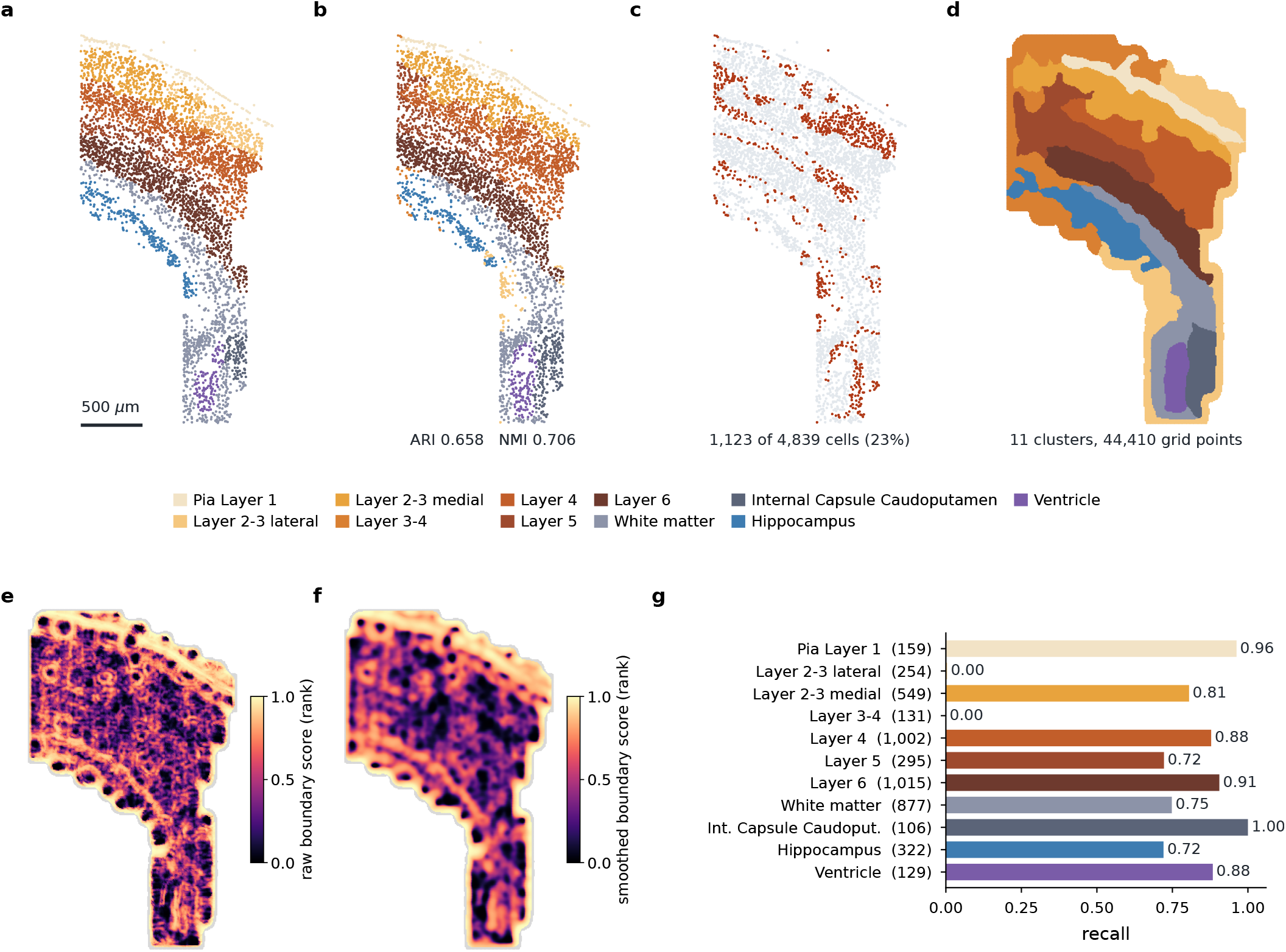
osmFISH mouse somatosensory cortex. Evaluation on the osmFISH cortical section (33 genes, 4,839 cells, 11 manually annotated cortical regions). **a**, Ground-truth annotation. **b**, Predicted domains, Hungarian-matched to the annotation (overall ARI 0.658, NMI 0.706). **c**, Misclassified cells (red; 1,123/4,839, 23.2%), concentrated at layer interfaces and in the two missed domains. **d**, Grid-level domains (11 clusters over 44,410 grid points), from which the cell-level labels are derived. **e**, Raw sheaf boundary score and **f**, the same field after Gaussian smoothing, each as its percentile rank among interior grid points (the tissue edge, grey, would otherwise set the scale); the score is highest along the pial surface, white matter and ventricle and lower at the layer interfaces. **g**, Per-domain recall across the 11 annotated regions (cell counts in parentheses); the internal capsule, pia, ventricle and Layer 6 are recovered at 0.88–1.00 and the other large domains at 0.72–0.88, whereas Layer 2–3 lateral and the thin 131-cell Layer 3–4 transitional zone are missed (recall 0).

SSAM reaches 0.729 here at the best of its grid by building domains from the composition of 74 de novo cell types (13 give 0.540); cortical layers are defined by which neuronal subtypes they contain, so a 74-dimensional composition vector carries laminar information that no low-dimensional projection of 33 genes holds. On a laminar cortex with a small panel the composition representation is better suited to the task.

### 2.6 Two independent cortical datasets

Both datasets above were used during development, so we added two annotated sections that were not: from the SpaceTx consortium [31], the Allen Institute prototype MERFISH section of mouse primary visual cortex (268 genes, 0.9 million molecules, 2,150 annotated cells in L1–L6b) and the Cold Spring Harbor BaristaSeq section [32] (81 genes, 0.32 million molecules, 4,432 cells in L1–L6 and white matter), from different laboratories and chemistries. Every method was tuned on each section over its own grid; spSheaf’s has 162 configurations, fewer than SSAM’s 287–288 (Table 1, Extended Data Fig. 2).

On BaristaSeq spSheaf reaches ARI 0.757 (0.749 at the local reference); SSAM 0.738, Points2Regions 0.733 and FICTURE 0.631, the three leading values within Points2Regions’ seed spread (0.685 ± 0.098). White matter, Layer 2/3 and Layer 4 are recovered at recall 0.94–0.95, Layers 5 and 6 at 0.79–0.80, and Layer 1 (32 cells) is missed: the sixth cluster is a *Synpr* -high patch within Layer 6 (201 cells, identical over four seeds; Extended Data Fig. 2b), and at *k* = 5 the patch merges into Layer 6 and the ARI rises to 0.782 (Extended Data Fig. 2b, last column; Section 2.7). On the Allen prototype spSheaf reaches 0.754; Points2Regions 0.684 (0.661 ± 0.052 over seeds), FICTURE 0.475, SSAM 0.410. On this section the coordinate reference determines the ranking: the local reference reaches 0.665, below Points2Regions, and the tissue reference 0.754, recovering L2/3 at recall 0.97 and L6b at 0.55, which the local reference splits or misses (Extended Data Fig. 2a). Over the 162 configurations the reference moves the mean ARI by 0.132 on Allen and 0.102 on BaristaSeq, more than any other parameter on Allen and second only to the spatial weight (0.141) on BaristaSeq (Extended Data Fig. 3), and the parameters interact: on Allen, switching the reference at the MERFISH scales gains 0.036 and widening the kernel then gains a further 0.176, whereas the same two changes under the local reference reach 0.643.

The reason for the dependence on the coordinate reference is unresolved. Molecule density does not explain it (osmFISH, the densest section, also prefers the tissue reference once each has its own search; MERFISH, at a similar density, does not), nor does boundary localisation (on MERFISH the tissue reference places the field slightly higher at annotated interfaces, 1.03 × against 0.93×, and still partitions worse). What differs on MERFISH is the width of the field after median normalisation: the tissue reference compresses it (99th percentile 2.34 against 2.78 times the median) and attenuates less. We do not infer a mechanism from a single dataset. Neither the reference nor *σ*_*s*_ has a label-free rule.

### 2.7 Choosing the number of domains without an annotation

spSheaf requires the number of domains *k* as an input. In the benchmarks above *k* is the annotated count, following the SDMBench protocol, and ARI against the annotation is maximal at that count on four of five MERFISH slices. For a section without an annotation, *k* must be chosen from the data. We evaluated the silhouette width of the fibers under the partition at each *k* — the standard internal criterion, computed on spSheaf’s own output — for *k* = 2–16 on all five MERFISH slices, against the annotated *k* = 8 (Extended Data Fig. 4). The silhouette is maximal at *k* = 10, 8, 9, 8 and 9 on the five slices, and on four of the five its value at *k* = 8 lies within 0.001 of its maximum. The curve is flat near its maximum, so the criterion identifies a narrow range of *k* containing the annotated value rather than the value itself; pooled over the five slices it is maximal at *k* = 9 (Section 2.8 examines the additional domain). The procedure we recommend is therefore to compute the silhouette over a range of *k* and to select within the interval over which it lies within 0.002 of its maximum, which on these sections contained the annotated count on four of five slices and was within two domains of it on the fifth. The BaristaSeq section shows the consequence of fixing *k* at the annotated count when one annotated domain is very small: Layer 1 holds 32 of 4,432 cells and is recovered by no method, spSheaf’s sixth cluster is spent on a *Synpr* - high patch within Layer 6, and at *k* = 5 the partition is the five remaining layers at ARI 0.782, against 0.757 at *k* = 6 (Extended Data Fig. 2b).

We also tested a criterion specific to the construction: the agreement (ARI) between the partitions obtained at two phases of the fiber lattice, offset by half a grid spacing, as a measure of whether the boundaries at a given *k* are supported by expression rather than by the discretisation. Read as an upper bound on *k* — the largest *k* at which the agreement remains within 5% of its value at *k* ≤3 — it recovers *K* on synthetic Voronoi tissue with known *K* (6–7 and 6–8 for *K* = 6; Section 4.13). On the MERFISH sections the same bound is 16, 15, 8, 12 and 9, and four of its ten lowest values coincide with drops in a *k*-means-seed control run at every *k*, indicating that on real sections the agreement is dominated by the multiplicity of *k*-means optima rather than by the lattice. This criterion is therefore not part of the recommended procedure; the full curves and the synthetic calibration are given in Extended Data Fig. 4.

### 2.8 Structure below the annotation

The silhouette’s preference for *k* = 9 can be examined on the sections by identifying the additional cluster. At *k* = 8, 9 and 10 on every slice, each cluster was assigned to the annotated domain holding the majority of its annotated grid points, a domain with two such clusters called *divided*, and the two clusters compared within the domain by share, purity, centroid offset and the difference of every fiber component, with a per-gene test that is descriptive only, since grid points are not independent samples (Section 4.8). Two divisions on different sections are the same division when their difference vectors correlate over the 135 genes. Supplementary Table 4 is the complete record, 26 divisions. Three kinds are set aside first: divisions already present at *k* = 8, because on four slices the thin periventricular PV receives no cluster and the cluster it does not use is spent elsewhere; six divisions of a bilateral domain into its hemispheres (centroids 1.2–1.3 mm apart on a 1.8 mm section), which are the spatial term, not expression; and two slivers of 7–9% of the PVH at purity below 0.45.

The remaining divisions are reproducible across sections. On the two posterior slices, the only two on which the BST is divided non-hemispherically, the annotated BST separates into two clusters of similar size, each at least 0.73 pure, with centroids offset along the same axis, and the difference vectors correlate at *r* = 0.84 across the two sections, with *Isl1, Esr1, Amigo2, Arhgap36, Coch, Gpr165, Gabrg1* and *Slc18a2* leading the smaller half on both; the division is unchanged from *k* = 8 to 10 (Fig. 4a,b). On the two anterior slices the MPN separates likewise (*r* = 0.88) into a *Calcr* /*Sgk1* /*Ermn* half and a *Npas1* /*Coch*/*Npy2r* /*Slc18a2* /*Esr1* /*Pgr* half at its medial edge (Fig. 4c,d), a division present at *k* = 8 whose smaller cluster is only half inside the annotated MPN, so it may mark the border rather than a compartment. The MPA divides on three slices in three unrelated ways (*r* from −0.28 to 0.43), twice with a vascular or mural signature (*Myh11, Lmod1, Rgs5, Pdgfra, Fn1*) at different locations, reflecting cellular composition rather than a subdomain. The two BST halves differ in the neuronal populations the dataset’s authors assigned to each cell [30], an annotation independent of the fibers the division was computed on. Assigning every annotated BST cell to the half of its nearest grid point, the smaller half holds 83% and 86% of the cells of inhibitory cluster I-13 on the two sections (odds ratio 7.3 and 8.2 against the other BST neurons, Fisher’s exact test, *q* < 10^−3^) and 75% and 74% of I-2, whereas I-23, E-4, I-9 and I-25 lie predominantly in the larger half (I-23: 14% and 25%); the per-cluster log odds correlate at *r* = 0.94 across the two sections and all eight clusters significant on either agree in sign (Fig. 4e). In the cells themselves, I-13 is distinguished from the other BST neurons by *Isl1, Arhgap36, Amigo2, Esr1, Slc18a2, Gpr165, Coch* and *Gabrg1*, the genes that lead the smaller half at grid level, and I-2 by *Crhbp, Brs3, Greb1, Prlr* and *Esr1* ; the grid-level signature of the division is the marker signature of the populations it encloses. On the hemispheric division of slice 0.14 the same clusters split with the opposite sign (*r* = −0.79 against slice 0.19), as a division by position rather than composition should. The smaller half is therefore the territory of two inhibitory populations of the source atlas, I-13 and I-2, within the annotated BST, a subdivision finer than the benchmark annotation. The I-13/I-2-enriched subdivision is consistent with a posterior BNST/striohypothalamic neuronal territory described in the source atlas and in a subsequent spatial re-analysis of it [30, 35], although the corresponding anatomical subnucleus cannot be assigned unambiguously.

**Fig. 4.**
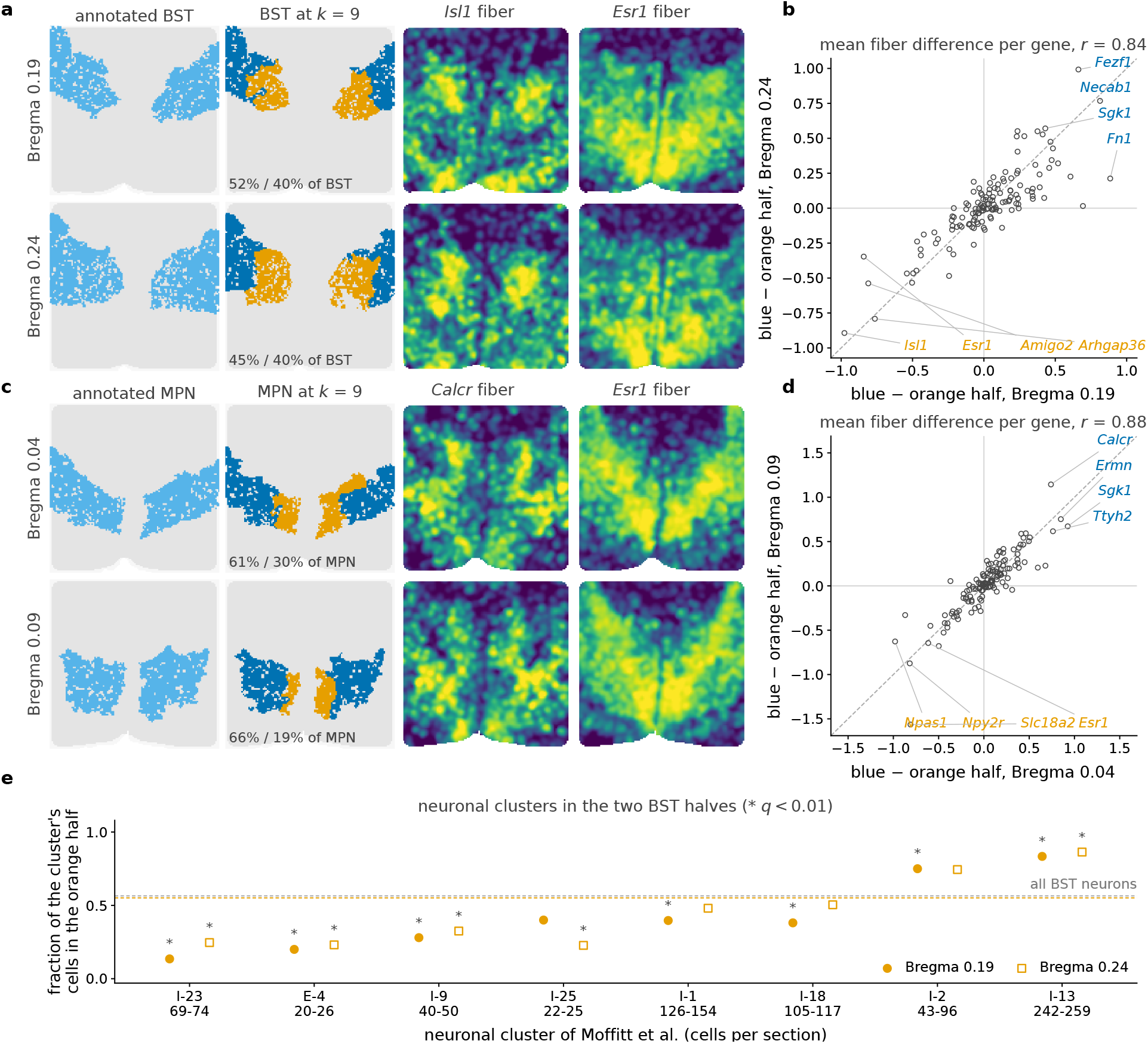
The extra domain at *k* = 9 is the same domain on adjacent sections. **a**, The annotated BST on the grid of slices 0.19 and 0.24; the two clusters at *k* = 9 whose majority annotation is BST, restricted to the domain (blue, the larger; orange, the smaller; shares printed); and the normalised *Isl1* and *Esr1* fibers over the section. **b**, The mean difference of every fiber component between the two halves on slice 0.19 against the same quantity on slice 0.24, one point per gene, *r* = 0.84 over 135 genes; the genes leading the difference on both are named (orange, higher in the smaller half; blue, in the larger). **c, d**, The same for the annotated MPN on slices 0.04 and 0.09, with the *Calcr* and *Esr1* fibers, *r* = 0.88. Shares, purities, offsets and the leading genes of every division, including those that do not repeat, are in Supplementary Table 4. **e**, The neuronal clusters of Moffitt et al. [30] in the two BST halves: for every cluster with at least 20 annotated BST cells, the fraction of its cells whose nearest grid point lies in the orange half, on slice 0.19 (filled) and 0.24 (open); dashed, the fraction of all BST neurons in that half on each slice; asterisks, *q* < 0.01 (Fisher’s exact test against the other BST neurons, Benjamini–Hochberg over clusters); cell counts per section under each cluster.

### 2.9 Beyond the benchmark datasets

Because the sheaf machinery acts in feature space, the same core applies to data of other kinds; none carries a curated spatial-domain ground truth, so evaluation is qualitative or by reproducibility. On the Stereo-seq E16.5 mouse embryo (*∼*552 million transcripts, 28,633 genes) and a Vizgen MERSCOPE whole coronal hemi-section (54.7 million molecules, 649 genes), spSheaf runs to completion after gene filtering and molecule subsampling and returns anatomically recognisable compartments (Extended Data Fig. 5); rigid registration of the MERSCOPE domains to the Allen Common Coordinate Framework gives ARI 0.264 against its ten major divisions, a value confounded with registration error. Through the feature-fiber variant it applies to three-dimensional subnuclear data — DNA seqFISH+ chromosome territories and sequential-immunofluorescence chromatin-mark zones [33] — and to multiplexed protein imaging [34] (Extended Data Figs. 6, 7 and 8), evaluated by cross-nucleus reproducibility and marker interpretation. On the carcinosarcoma section the inferred domains partition an epithelial–mesenchymal continuum with one discrete mesenchymal compartment; across that compartment’s edge the EMT score changes by 0.71 over about five grid spacings in replicate 1 (a logistic width of one spacing, 150 px, at a kernel bandwidth of 300 px, i.e. at the resolution of the fibers) and by 0.16 in replicate 2, with the disagreement field at the edge 1.05× and 0.95× its interior value (Extended Data Fig. 8g,h), so the compartment edge is an interface in expression at the resolution measured in one section and a weak gradient in the other. Fibers, stalks, restriction maps and the disagreement score are unchanged across these; what varies is the fiber contents and the spatial discretisation.

## 3 Discussion

spSheaf infers spatial domains from raw molecule coordinates without cell segmentation. Its central quantity is the sheaf disagreement of a cellular sheaf over a spatial graph: the residual incompatibility of the local expression coordinates carried by neighbouring stalks once those stalks have been aligned. Transitions between expression programs raise it, and attenuating the regions where it is high yields the partition.

### Contribution of the sheaf construction

The ablation quantifies each component. Weighting by a smooth field of local expression change accounts for most of the improvement over no weighting: the kernel baseline, which compares raw neighbouring fibers with no alignment and no local basis, is 0.046 ARI behind the method, against 0.102 for no weighting. The remaining 0.046 arises from two choices: comparing coordinates transported by the restriction maps rather than fibers (+0.010 over identity maps), and referencing each stalk’s coordinates to its own neighbourhood (+0.040). The ablation is at the configuration selected on this benchmark, which uses the local reference; at the method’s default, the tissue reference, the score is 0.062 above no weighting and 0.006 above the kernel baseline on these five slices; the local reference, selected on this benchmark, is what makes the alignment count: it removes each stalk’s own neighbourhood mean before transport, so the transported difference measures the change of program across the edge and not the shared offset of both neighbourhoods from the section mean (Section 4.5); on the two SpaceTx sections the weighting itself contributes 0.012–0.024 ARI and no edge score separates from the kernel baseline (Supplementary Fig. 3). Scores that compare the subspaces themselves are no better than a random field at partitioning (0.491–0.495 against 0.474), although they localise annotated boundaries best of the nine scores (57 *µ*m against 160 *µ*m for the method). These observations are consistent: a subspace score responds within a narrow band at the interface, a transported-coordinate score across the full transition between programs, and the full transition is what must be attenuated for the clustering to separate the two programs. The sheaf construction is the setting in which the comparison of aligned coordinates, rather than of subspaces, is defined, and the alignment it performs is not degenerate (median largest principal angle between adjacent stalks 46^°^; Section 4.5). Its gain over kernel smoothing is specific to tissue whose domains are distinct programs, +0.046 ARI on the hypothalamus; on laminar cortex, where neighbouring stalks lie on the same program, the aligned and the raw comparison agree and the score is neutral.

### Why discontinuities rather than the spectrum

A natural alternative is to assemble the sheaf Laplacian and use its low-frequency eigenvectors [27]. We did not, for a structural reason: those eigenvectors are by construction the smoothest modes over the graph, whereas an interface is a discontinuity, so such a method emphasises the quantity that is not to be localised; manifold embeddings (Isomap, UMAP) smooth for the same reason. All three were tried during development and not carried further; no systematic comparison is reported. Vector diffusion maps [28] and ddHodge [29] build the same local bases and SVD alignments; the differences are the base space (a spatial grid rather than a graph on the data manifold), the coordinates (data projections rather than diffusion coordinates) and the downstream operation (a residual, which measures discontinuity, rather than an embedding, which smooths). The sheaf Laplacian assembled from our restriction maps would be the connection Laplacian restricted to the spatial graph.

### Scope

spSheaf assumes that adjacent domains differ in the geometry of local expression, and the osmFISH section shows the limit of that assumption: on a 33-gene laminar cortex two of eleven layers are not recovered at any setting, and a cell-type-composition representation of the same data does better, as SSAM’s 74 de novo types show. The two SpaceTx sections, also laminar, show the same distinction from the ablation side: the full pipeline leads or ties the competitors on both, but the sheaf term matches kernel smoothing there (Supplementary Fig. 3), whereas on the hypothalamus, whose nuclei are distinct programs with sharp interfaces, it contributes +0.102 ARI. The disagreement score measures a change of local expression program between neighbouring stalks; along a gradient of composition over a shared set of cell types, neighbouring stalks lie on the same program and the score carries no interface. The method therefore suits tissue whose domains are distinct expression programs with sharp interfaces — nuclei, compartments, zonation — and a panel that resolves those programs; on tissue organised as a compositional gradient, such as cortical layers, the pipeline remains competitive and the sheaf term is neutral. Its evidence base is four annotated mouse-brain datasets and three segmentation-free comparators run on identical inputs; segmentation-based methods are represented only through the published benchmark, and human tissue and non-neural tissue are untested. The method is designed for tissue organisation, not for single-cell profiles: by operating on a grid of kernel fibers it does not inherit the segmentation-dependent misassignment documented across platforms [14], but neither does it produce the per-cell expression profiles that cell-type annotation and ligand–receptor inference require. The applications beyond those sections demonstrate portability across dimension and modality, not accuracy.

### Limitations

The number of domains *k* is chosen by the silhouette procedure of Section 2.7, which identifies a range of *k* rather than a value — within two domains of the annotation on all five MERFISH sections — and which, on synthetic tissue whose domains differ only subtly, under-counts. Two further settings have no label-free rule: the coordinate reference, whose default, the tissue reference, is the better of the two on three of the four annotated datasets and the worse by 0.02 ARI on the MERFISH benchmark, and which accounts for more of the grid’s variation than any other parameter on both SpaceTx sections; and the smoothing radius *σ*_*s*_, whose selected value ranges from 58.5 to 180 *µ*m across the four datasets; a rule for these two from the data alone is the most immediate practical improvement. Tuned on the annotation it is scored against, spSheaf leads on two of four sections and ties on a third; untuned, with every parameter carried over from MERFISH, it trails each section’s best tuned competitor by 0.03–0.15 ARI (Extended Data Table 1), which is the expected performance without tuning. Cost is linear in grid points and edges, about 30 s per MERFISH section on one CPU core at 14,286 grid points; competitors’ runtimes were not measured under matched conditions and are not compared. Compartment identities on the datasets without ground truth are marker interpretations that require expert confirmation; the BST division of Section 2.8 is anchored to the cell populations of the source atlas, but its anatomical name is not.

## 4 Methods

### 4.1 Data description

We applied spSheaf to imaging- and sequencing-based spatial datasets spanning RNA, DNA, and protein. The **Moffitt et al. MERFISH dataset** [30] profiles the mouse hypothalamic preoptic region (135 genes, 20.5 million molecules in the full export, 5 annotated slices with 5,488–5,926 annotated cells each and 8 anatomical domains); processed molecule tables were prepared by Y. Imoto. Each section is imaged in seven focal planes 1.5 *µ*m apart; every method, spSheaf included, reads every molecule of the section — selected by its bregma coordinate, all planes projected onto the section plane, the molecules of unannotated cells included — which is 1.8–2.0 million molecules and 6,100–6,600 cells per section, and is scored on the 5,488–5,926 annotated cells (Section 4.11). The **osmFISH mouse somatosensory cortex** [6] comprises a 33-gene panel over 4,839 annotated cells in 11 regions; the raw molecule table read by every method carries 1,976,659 molecules under 35 gene labels (the panel plus *Klk6* and *Lum*, which the annotation’s cell matrix omits) from all 39 hybridisation rounds, including the six the original study flagged as low quality (174,070 molecules); the same table, with two legacy labels renamed, is the competitors’ input. Two **SpaceTx** mouse primary visual cortex sections [31] were taken from the consortium’s public release, each as a molecule table (`spot table`, columns `xc, yc, gene`) and a segmented-cell table with a per-cell `layer` annotation, both in micrometres: the **Allen Institute prototype MERFISH** section (268 genes, 900,678 molecules, 2,150 annotated cells in L1–L6b, of which L1 has 9, over 1.3 × 1.1 mm) and the **Baris-taSeq** section [32] from Cold Spring Harbor Laboratory (81 genes, 319,358 molecules, 4,432 cells in L1–L6 and white matter, of which L1 has 32, over 0.8 × 1.3 mm). Neither section was used in development. The **Stereo-seq E16.5 mouse embryo** [9] contains *∼*552 million detections over 28,633 genes. The **Vizgen MERSCOPE mouse-brain receptor showcase** is a whole coronal hemisection (54.7 million molecules, 649 genes, 7 focal planes). The **Takei et al. DNA seqFISH+ and sequential-immunofluorescence** data [33] provide, per single mouse embryonic stem-cell nucleus, 3,600 genomic-locus coordinates (xy 103 nm, z 250 nm) and pervoxel intensities of 21 chromatin marks. The **PECAb carcinosarcoma** [34] data provide a 35-antibody protein panel and a companion 220-gene panel over 7,657 cells. Accessions are listed under Data availability.

### 4.2 Problem formulation

Let 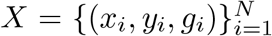, with (*x*_*i*_, *y*_*i*_) *∈* ℝ^2^ the coordinates of the *i*-th molecule and *g*_*i*_ *∈* {1, …, *G*} its gene identity. The tissue is partitioned into *k* spatial domains Ω_1_, …, Ω_*k*_ *⊂* ℝ^2^ with distinct expression programs; the goal is to recover this partition from *X* without cell segmentation.

### 4.3 Expression fiber construction

A regular grid 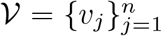 with spacing *δ* is laid over the tissue. At each grid point *v*, the expression fiber *f*_*v*_ *∈* ℝ^*G*^ is

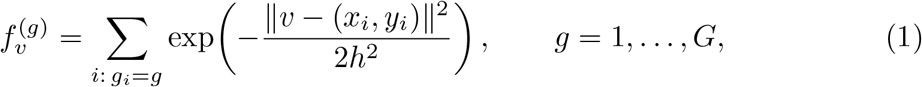

with bandwidth *h* and search radius 3*h*; grid points with total weighted count below a threshold are discarded. Fibers are normalised to proportions and log-transformed,

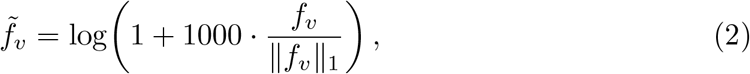

with 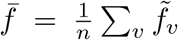 the mean fiber of the section, used by the global PCA of the clustering step and by the optional tissue reference below.

### 4.4 Continuous feature fibers and the N-dimensional grid

When each point carries a continuous feature vector *ϕ*_*i*_ *∈* ℝ^*P*^ (per-locus chromosome intensities, per-voxel chromatin-mark intensities, per-cell antibody or gene panels) rather than a categorical gene identity, the fiber is the kernel-weighted mean

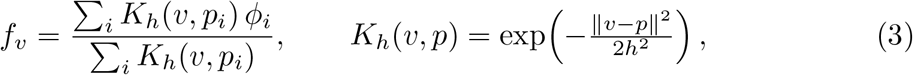

subsequently standardised per feature. For volumetric data the grid and kernel extend to ℝ^3^ with independent axial spacing and bandwidth (*δ*_*z*_, *h*_*z*_), implemented as a bandwidth-scaled isotropic kernel; the graph, stalks, restriction maps, disagreement, and clustering are unchanged, and boundary smoothing uses a *k*-d-tree Gaussian in three dimensions. This variant produces every non-transcriptomic and 3D result in this study without altering the geometric core.

### 4.5 Spatial graph, stalks, restriction maps, and disagreement

A symmetric *k*_graph_-nearest-neighbour graph *G* = (*V, ℰ*) is built on the grid (*k*_graph_ = 6). For each vertex *v*, let *N* (*v*) denote its *k*_nbrs_ nearest grid points (*v* itself included) and 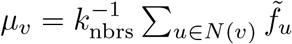 their mean. The matrix 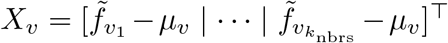 collects the *locally* centred neighbour fibers, with singular value decomposition 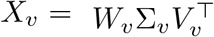 the stalk basis is 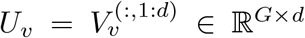 The stalk is thus the leading eigenbasis of the local covariance and is invariant to any global shift of the fibers. The local coordinates of *v* are its own fiber’s scores in that basis, referenced to the same neighbourhood mean, 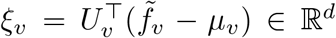 invariant to any shift of the fibers and independent of what else the tissue section contains. Referencing *ξ* to the tissue mean instead, 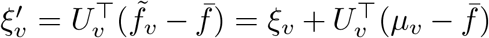, is kept as an option (*tissue reference*). It adds to the transported difference on every edge *e* = (*u, v*) the exact term 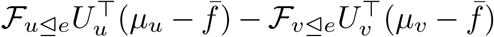, which does not vanish when the stalks are aligned: for *U*_*u*_ = *U*_*v*_ = *U* it reduces to *U* ^*⊤*^(*µ*_*u*_ − *µ*_*v*_), the projected difference of the two neighbourhood means. The tissue reference therefore re-introduces a change of mean expression program into the score, and where the stalks are differently oriented it also carries their misalignment applied to each neighbourhood’s offset from the tissue mean; it makes the score depend on the section’s composition, and whether it adds signal or variance is an empirical question, answered per dataset and not in general (Sections 2.4, 2.6). The default is the tissue reference; the local reference is retained as an option and is included in the tuning grid (Section 4.14), where it is the value selected on the MERFISH benchmark and the tissue reference the value selected on the three other annotated sections. A uniform stalk dimension *d* is used throughout; on the three single sections ARI is insensitive to *d* over the range swept, 4 ≤ *d* ≤ 20 (Extended Data Fig. 3); on MERFISH *d* = 6 was fixed during development. For each edge *e* = (*u, v*), the overlap 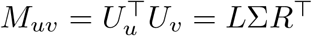 yields restriction maps *ℱ*_*u⊴e*_ = *L*^*⊤*^ and ℱ_*v⊴e*_ = *R*^*⊤*^, whose singular values are cosines of the principal angles between the stalks [36]. The sheaf disagreement is

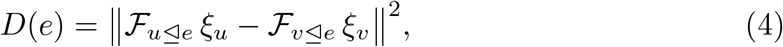

the residual mismatch after optimal subspace alignment. Unlike the Grassmann distance, which depends only on the principal angles, *D*(*e*) also depends on the projected expression profiles *ξ*_*u*_, *ξ*_*v*_: two stalks spanning the same subspace can still disagree. The score is therefore properly read as an alignment-adjusted disagreement of local coordinates, not as a pure measure of subspace rotation. Adjacent stalks are not near-coincident even though neighbouring vertices share most of their *k*_nbrs_ points: on the five MERFISH slices at the reference configuration the largest principal angle between *U*_*u*_ and *U*_*v*_ has median 46^°^ over the 50,100–50,800 edges of each slice (90th percentile 76^°^; mean over the *d* = 6 angles 16^°^), against 75–83^°^ and 40–46^°^ for stalks fitted to random non-local neighbourhoods of the same size, and the aligning rotation 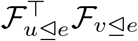 is at median Frobenius distance 2.9 from the identity (maximum 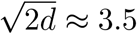 experiments/merfish/stalk angles.py).

### 4.6 Boundary score, clustering, and cleanup

The per-vertex boundary score is 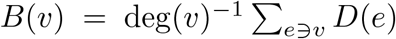 (mean aggregation for robustness to outlier edges), rasterised to the grid, convolved with a Gaussian of standard deviation *σ*_*s*_, and resampled at the grid points as 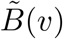. For clustering, a global PCA embedding 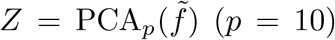 is standardised; the exponential interior weight is

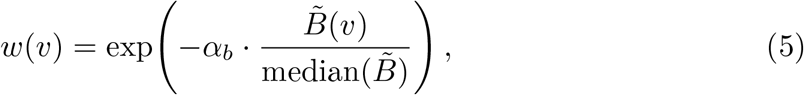

applied row-wise to 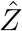 standardised spatial coordinates *S* are concatenated as 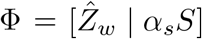 and clustered by *k*-means with *k*_clusters_ clusters and multiple initialisations. Interior points cluster by expression; boundary points, whose features are attenuated, defer to spatial proximity. Finally, connected fragments smaller than a fraction *τ* of all grid points (at least three vertices), excluding each label’s largest component, take the label of an adjacent non-fragment vertex, in up to five passes of breadth-first search on *G*.

### 4.7 Multi-slice alignment

For serial slices, per-gene expression-weighted centroids 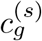 define landmarks; interslice shift vectors 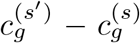 are computed, a radius graph on the shifts is built, connected components are extracted by breadth-first search, and the dominant component gives the consensus translation (direct-to-reference, no chaining). Predicted labels of adjacent slices are then matched by the Hungarian algorithm on their co-occurrence matrix over inter-slice edges within 45 *µ*m (Supplementary Fig. 2).

### 4.8 Evaluation and baselines

Where ground truth exists, we report the Adjusted Rand Index (ARI) and Normalised Mutual Information (NMI), with predicted labels remapped by the Hungarian algorithm for visualisation. Following the SDMBench protocol [24], the pipeline is run on the raw molecule table without access to the annotation and grid predictions are mapped to cell centroids by nearest-grid-point assignment before scoring at cell level; ground-truth labels are never injected before the run. Baseline numbers for the SDMBench comparison are taken from the source data of Yuan et al. [24]; the three segmentation-free comparators were run directly on the identical molecule tables (Section 4.11). The naive baselines of Section 2.5 are *k*-means into the annotated number of groups on the standardised cell centroids, the standardised cell-by-gene matrix, and the two concatenated, over five seeds of ten initialisations each; on the SpaceTx sections (Section 2.6) the expression input is log-normalised and reduced to 10 principal components before standardisation, over 20 seeds. The *k*-sweeps of Section 2.7 vary *k*_clusters_ with all else fixed: one over 4–16 scored at grid level by ARI and NMI and by the silhouette width of the normalised fibers and the within-domain fraction of spatial-graph edges, and one over 2–16 on all five slices carrying grid-phase stability, its seed control and the full-population silhouette (Section 4.13; Extended Data Fig. 4). The division analysis at *k* = 8–10 assigns every cluster to the annotated domain holding the majority of its annotated grid points, calls a domain divided when two clusters share it, and compares the two largest by the mean of every normalised fiber component within the annotated domain, with a per-gene Mann–Whitney test, Benjamini–Hochberg corrected, on the same points (experiments/merfish/k split.py; Fig. 4, Supplementary Table 4). The composition of the two BST halves (Fig. 4e) uses the cell-level neuronal cluster labels of Moffitt et al. [30]: every annotated BST cell is assigned to the cluster of its nearest grid point (median distance 6 *µ*m), cells on clusters other than the two halves are dropped (9–14%), and for each neuronal cluster with at least 20 cells the odds of lying in the smaller half against all other BST neurons are tested by Fisher’s exact test with Benjamini–Hochberg correction over clusters; cluster marker genes are computed from the per-cell molecule counts normalised to counts per thousand (experiments/merfish/k split celltypes.py). Where no spatial-domain ground truth exists we report unsupervised metrics (within-domain edge fraction, silhouette), data-driven marker genes, and, for the nuclear data, cross-nucleus and cross-replicate reproducibility.

### 4.9 Edge-score ablation and direct boundary evaluation

Every arm of Fig. 2d replaces only the edge score *D*(*e*); the fibers, grid, graph, stalks, weighting function, clustering and cleanup are byte-identical across arms, and all arms share the same five *k*-means seeds on each of the five slices. Writing 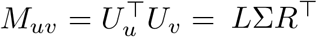 and 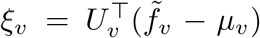, the arms are: *sheaf (local ref*.*), ∥L*^*⊤*^*ξ*_*u*_ − *R*^*⊤*^*ξ*_*v*_*∥*^2^, the full construction; *sheaf (tissue ref*.), the same with 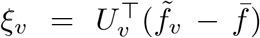 the optional tissue reference; *identity*, 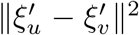 with the tissue-referenced coordinates 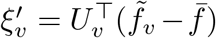 omitting the alignment; *neighbour*, 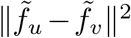, a kernel baseline with no sheaf structure; *grassmann*, the Grassmann distance 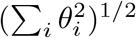 from the principal angles *θ*_*i*_ = arccos *σ*_*i*_; *projection*, 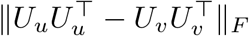 ; and *procrustes*, the squared residual 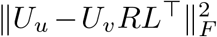 of the optimal orthogonal alignment. The three null controls are *shuffled* (the sheaf field permuted uniformly at random across vertices, preserving its marginal distribution and destroying its spatial arrangement), *random* (an independent uniform random field), and *none* (*α*_*b*_ = 0, i.e. no weighting).

For the direct boundary evaluation (Section 2.4), ground-truth boundary vertices are those grid points adjacent in the spatial graph to a vertex carrying a different annotated label, after mapping the cell annotation to the grid by nearest-molecule assignment; the resulting positive rate is 14.5–16.4% per section. Each candidate field is thresholded over a sweep of *selection fractions* (the top *q* of vertices for *q* on a fixed grid) rather than over score quantiles, so that fields with differently shaped score distributions are compared at matched selection sizes; average precision is the standard step-interpolated area under the resulting precision–recall curve. Localisation is reported as the mean distance from each predicted boundary vertex (the top *q* of vertices, *q* equal to the true positive rate) to the nearest true boundary vertex, and boundary-F1 at a 15 *µ*m tolerance. This evaluation is performed at grid level, where the field is defined, whereas the ARI and NMI of Section 2.2 are scored at cell level; ground-truth boundaries inherit the coarseness of the annotation. All arms use the MERFISH *α*_*b*_, *σ*_*s*_ and graph, and their ARI and NMI are scored at grid level.

A controlled construction with two regions of identical mean expression and identical neighbour-distance distributions but fibers in orthogonal subspaces reproduces the ordering of Section 2.4 — subspace geometry localises the interface, the coordinate score responds over the whole transition — and is kept in the repository (`make method-validation`) rather than in the paper.

### 4.10 osmFISH coordinate registration

The osmFISH benchmark annotation and the raw molecule table are not in the same frame: the cell coordinates of the benchmark AnnData follow the published loom file, whereas the molecule table is in the raw imaging frame, and the two are flipped on both axes and translated relative to each other. The transform was recovered without using the domain labels, by ranking candidate orientations on how well the gene composition of molecules within a radius of a cell centroid reproduces that cell’s own count vector; the chosen transform reaches a mean per-cell composition correlation of 0.83 within 60 px (3.9 *µ*m) of a cell centroid against 0.07 for the unflipped alignment. Molecule coordinates are in camera pixels at 15.3846 px/*µ*m and are converted to micrometres before rasterisation. Both corrections apply to every method in the comparison. The spSheaf evaluation registers the annotation onto its molecule grid by the same label-free criterion (mean composition correlation within a cell radius, 0.53 at the recovered translation against 0.17 for centroid matching); the two differ by (109, −74) *µ*m, and centroid matching scores the same partition at ARI 0.538.

### 4.11 Segmentation-free comparison protocol

All three comparators were run with their published implementations on the identical raw molecule tables (SSAM’s `find_domains` sweep on MERFISH through a line-for-line re-implementation, checked identical on slice 0.19). A comparator qualifies if it reads the molecule table and returns a spatial partition of the section; segmentation-free tools that stop at a per-pixel cell-type map, such as Sainsc [17], were not included, because a domain stage added by us would be scored in their place, and SSAM represents that family with its authors’ own domain-inference step. On MERFISH that table is every molecule of the physical section, selected by its bregma coordinate, in all seven focal planes (Section 4.1); on osmFISH every method is scored in the registered frame of Section 4.10; on the two SpaceTx sections the molecule and cell tables share one frame, confirmed by the same label-free criterion (composition correlation 0.94 at centroid alignment and 0.97 after an 8 *µ*m shift on the Allen prototype; 0.89 and 0.90 at 4 *µ*m on BaristaSeq), and the recovered translation is applied to every method’s cell placement. Metrics are a reimplementation of the SDMBench definitions [24]. The domain metrics require one label per annotated cell: grid- or pixel-valued output (SSAM’s `inferred_domains`, the tissue-domain field over all pixels) was sampled at each cell centroid; molecule-labelled output (Points2Regions) was assigned by majority vote over the 30 molecules nearest the centroid; cells left unassigned or masked as background were filled from the nearest labelled cell. FICTURE’s pixel-level output (top-3 factors and posterior probabilities per pixel) was converted to domain labels three ways — the modal top-1 factor of pixels within 30 *µ*m; *k*-means on the mean factor-probability vector of those pixels; and the same with standardised spatial coordinates appended at *α*_*s*_ = 0.5 — and the mapping is part of its configuration.

*SSAM* followed the authors’ notebooks (`HiDiHlabs/ssam example`; MERFISH settings on the SpaceTx sections). Its tissue domains are derived from its cell-type map, and the published parameters were tuned against SSAM 1.0: in release 1.1.3 the same call at Louvain resolution 0.15 with the default outlier filter leaves only 5–7 cell types per MERFISH section, against the 68 (curated to 50) of the published analysis, and on the SpaceTx sections it returns no cell types at all. We therefore restored the published `sctransform` normalisation (R 4.3.3, `sctransform` 0.4.1), disabled the outlier filter, and treated the resolution as a hyperparameter, running it at 4 and 8 on MER-FISH (39–44 and 61–63 cell types), at 0.15, 4 and 8 on osmFISH (13, 51, 74) and at 1, 4 and 8 on the SpaceTx sections (10–66), alongside the built-in normalisation — at the published resolution 0.15 with the filter on for MERFISH (5–7 types) and osmFISH (12), and at resolution 1 with the filter off on the SpaceTx sections (13 and 20), where the published recipe returned none. Each clustering is carried through a `find_domains` sweep: on MERFISH `n_clusters` *∈* {8, 12, 20} × `merge_thres` *∈* {0.6, 0.7, 0.8, 0.9} × `norm_thres` *∈* { 1000, 4000} with `merge_remote` fixed (24 per clustering, 72 per section); on osmFISH `n_clusters` *∈* { 11, 15, 20} × `merge_thres` *∈* { 0.6, 0.7, 0.8} × `merge_remote` × `norm_thres` *∈* { 1500, 4500} (36 per clustering, 144 in all); on the SpaceTx sections `n_clusters` *∈* { 6, 12, 20} `merge_thres` 0.6–0.9 × `merge_remote` × `norm_thres` *∈* { 300, 600, 1000} (71–72 per clustering, 287–288 in all; on the Allen prototype the published `norm_thres` of 4000 leaves no domain). The published recipe is never selected: on MERFISH the held-out rule picks resolution 4 or 8 on every fold, raising SSAM’s held-out mean from 0.131 to 0.343; on osmFISH the three resolutions give 0.540, 0.675 and 0.729 against 0.566 for the built-in normalisation; on the Allen prototype the built-in normalisation stays best (0.410) and on BaristaSeq `sctransform` at resolution 1 raises SSAM from 0.703 to 0.738. Two limitations remain: `find_domains` labels connected components, so the number of output domains cannot be fixed in advance (2 to 54 across the MERFISH sweep, 1 to 92 on the Allen proto-type); and the published SSAM analyses curate clusters by hand, a step the resolution sweep stands in for without reproducing.

*Points2Regions* was grid-searched over `pixel_width` × `pixel_smoothing` with the cluster count fixed to the annotated number of domains: 45 configurations per MERFISH slice, 32 on osmFISH, and {5, 10, 20, 40} *µ*m × {1, 2, 4, 6, 9, 15} on the SpaceTx sections (24); five seeds were run at each selected setting. *FICTURE* was run through its `run_together` pipeline (v0.0.5.7) at factor counts {8, 12, 20} (MER-FISH), {11, 15, 20} (osmFISH) and {6, 12, 20} (SpaceTx), each at training widths 12 and 18 *µ*m and with the three mappings above (18 configurations per section; its seeded mappings were also run at five seeds, 66 runs). It scores higher with 20 factors aggregated than with the factor count set to *k*, consistent with its design, in which factors are cell-type-like programs and domains their local composition.

*spSheaf* ‘s grids are 27 configurations per coordinate reference (54 in all) on MER-FISH (Section 4.14); on osmFISH 108 per coordinate reference (grid spacing *∈* {100, 150, 200} px, bandwidth *∈* {300, 500, 700} px, smoothing radius *∈* {300, 500, 900} px, *d ∈* { 6, 9}, *α*_*s*_ *∈* {0.3, 0.5 }, *α*_*b*_ = 0.5; 216 in all, plus 24 extending *d* to {12, 16, 20} over bandwidth {300, 500} px, radius {300, 500} px and both spatial weights under the local reference), with argmax 0.658 over the 108-configuration local grid (the configuration of Fig. 3) and 0.673 under the tissue reference (bandwidth 300 px, radius 900 px, *d* = 9, *α*_*s*_ = 0.3; the configuration of Supplementary Table 3); and on each SpaceTx section 162 (coordinate reference *∈* {local, tissue}, *h ∈* {25, 40, 60} *µ*m, *α*_*s*_ *∈* {0.25, 0.5, 0.75}, *σ*_*s*_ *∈* {60, 120, 180} *µ*m, *d ∈* {4, 6, 9}, at *δ* = 10 *µ*m and *α*_*b*_ = 0.5), below SSAM’s 287–288 and above FICTURE’s 18 and Points2Regions’ 24. Table 1 reports the argmax of each grid restricted to each coordinate reference; the selected settings are in Supplementary Table 3.

### 4.12 Matched model selection

Every method was run over a parameter grid on every MERFISH slice (Points2Regions 45 configurations, SSAM 72, FICTURE 18, spSheaf 27 per coordinate reference, 54 in all; Section 4.14), and every configuration’s score on every slice was retained. Three rules are then applied identically to all four methods: the per-slice argmax; the one configuration with the highest mean ARI over the five applied to all five (the rule of Table 1, comparable to the single sections); and leave-one-slice-out, where for each held-out slice the configuration maximising mean ARI on the remaining four is selected and its score on the held-out slice is reported (spSheaf: held-out mean 0.552, four distinct fold winners, each within 0.008 of the development configuration on its training slices). One rule governs what counts as a configuration: a single run at a fixed seed — one run per (pixel width, smoothing) for Points2Regions, per `find_domains` setting for SSAM, per configuration at *k*-means seed 42 for spSheaf (on MERFISH Table 1 reports the 20-seed mean of Supplementary Table 1; the seed-42 run scores 0.571 / 0.655). FICTURE is the only method with seed replicates and is taken at its first seed: averaging its seeds would score it at an expected value while every other method sits at one draw, and the best of all its runs would hand it a wider search than any other method was given (both alternatives are recorded in the released comparison table). osmFISH and the two SpaceTx sections admit no held-out slice, so on those the in-sample argmax is the only rule available and every method is under it (make matched-selection).

### 4.13 Synthetic tissue for the choice of k

A benchmark annotation cannot settle whether a criterion finds the right number of domains, because the annotated count is a granularity an expert chose. The criteria of Section 2.7 are therefore calibrated on constructions in which the count is known by construction.

Each construction places *K* seed points at random in a square field and assigns every molecule to the Voronoi cell of its nearest seed, so domains are contiguous and irregular rather than banded, and the spatial term cannot recover them for free. Each domain draws its genes from its own multinomial over 60 genes, obtained by softmax of a random unit direction scaled by a separation parameter; 2.2 × 10^5^ molecules are drawn in total, giving a per-fiber molecule count of the order of the real sections at the grid spacing used (*δ* = 12, *h* = 22, *σ*_*s*_ = 50, *d* = 6, *α*_*s*_ = 0.5). The resulting molecule table is passed to the same pipeline as the real data, and the generating domain of a grid point is its nearest seed.

Two designs are used. In the *flat* design the *K* programs are independent directions, so *K* is unambiguous and a criterion that cannot find it there will not find it anywhere. In the *nested* design two pairs of domains are made siblings — the second of each pair is the first plus 0.22 of an independent direction, renormalised — so *K, K*−1 and *K* − 2 are all defensible readings of the same tissue, which is the synthetic analogue of an annotation’s granularity being a choice. Each design is run at two independent draws of seeds, programs and molecules.

Grid-phase stability compares a run at the default lattice with runs offset by half a grid spacing in both axes. The two runs have different vertices, so the shifted labels are carried to the default lattice by nearest neighbour before the ARI is taken; the comparison is therefore between partitions of the same points. The *k*-means-seed agreement is computed alongside as a control at every *k*, and is reported because on the real sections it is what distinguishes a boundary moved by the lattice from a clustering with two near-degenerate optima. The silhouette is computed on the normalised fibers over every grid point: on a 4,000-point subsample it differed from the full-population value by up to 0.014, which is enough to move the argmax of a curve whose top is flat to within 0.001. The procedure and the constructions are released as experiments/synthetic/k selection.py and, for the real sections, `experiments/merfish/k selection.py.`

### 4.14 Hyperparameters

Supplementary Table 2 gives the MERFISH configuration. *k*_graph_ and *k*_nbrs_ were never tuned; *δ, h* and *d* were set on the MERFISH slices during development; (*σ*_*s*_, *α*_*b*_, *α*_*s*_) are the argmax over the five slices of a 54-configuration grid (*σ*_*s*_ *∈* {30, 60, 120} *µ*m, *α*_*b*_ *∈* {0.25, 0.5, 1}, *α*_*s*_ *∈* {0.3, 0.5, 0.7}, at each coordinate reference; `make merfish-loo-cv`), mean ARI 0.571 against 0.570 for *σ*_*s*_ = 120 *µ*m. This grid is spSheaf’s search on MERFISH under all three selection rules of Section 4.12: its perslice argmax is 0.584 and its leave-one-slice-out value 0.552 ± 0.052, with the selected configuration changing across folds (four distinct winners, all at the local reference). The tissue reference, the method’s default, reaches 0.548 under the one-configuration rule at *σ*_*s*_ = 120 *µ*m and 0.540 held out, with a seed spread of ±0.006 against ±0.000 for the local reference. With everything else identical the local reference scores 0.576 against 0.536 for the tissue reference over five slices and five seeds, ahead by 0.03–0.10 on three slices and within 0.004 on the other two (Fig. 2d).

Supplementary Table 3 lists the per-dataset settings. On the three single annotated sections they are the argmax of the grids of Section 4.11, selected on the section’s annotation like every competitor’s; *k*_graph_, *k*_nbrs_ and *α*_*b*_ are the MERFISH values throughout. On the datasets without annotation (Supplementary Table 3, lower rows) the scales are set by hand from the platform’s molecule density and physical extent, and nothing is selected.

### 4.15 Implementation and availability

spSheaf is implemented as the Python package `spsheaf`, depending only on standard scientific Python libraries (NumPy, SciPy, scikit-learn, pandas), and is installable via `pip install -e` .. All hyperparameters are exposed as command-line arguments in per-experiment entry-point scripts.

### 4.16 Statistics and reproducibility

No statistical method predetermined sample size; all 5 MERFISH slices, all osmFISH cells, all annotated cells of the two SpaceTx sections, and all available nuclei were used. No annotated cells or sections were excluded; the two whole-section demonstrations use the gene filtering and molecule subsampling of Supplementary Table 3. Experiments were not randomised and investigators were not blinded, as no experimental intervention was involved. ARI and NMI are reported per slice and as means; standard deviations over seeds or folds are population standard deviations (ddof = 0). On osmFISH the reported run is seed 42 (the rule of Section 4.12); over 20 initialisations at the same configuration ARI is 0.648 ±0.012 (median 0.653, range 0.620–0.662), against ±0.001 on MERFISH. The edge-score ablation reports means over 5 slices × 1*k*-means seeds, all arms sharing the same seeds and fibers. Large language models were used as coding assistants in preparing the initial runs of the competitor methods (Section 4.11) and in editing the text; all code, analyses and results were checked and are reproduced by the released repository, and the authors take full responsibility for the content.

## Data availability

The Moffitt et al. MERFISH data are available on Dryad (doi:10.5061/dryad.8t8s248) [30]; processed molecule tables were prepared by Y. Imoto and are available on reasonable request. The osmFISH data [6] are available from the original study (http://linnarssonlab.org/osmFISH); the MER-FISH and osmFISH domain annotations are the processed AnnData files released with SDMBench [24]. The SpaceTx Allen-prototype MERFISH and BaristaSeq molecule and cell tables are available from the consortium’s data page (https://spacetx.github.io/data.html) [31, 32]. The Stereo-seq E16.5 embryo data are in the MOSTA database (https://db.cngb.org/stomics/mosta, STDS0000058) [9]. The Vizgen MERSCOPE mouse-brain receptor showcase is available from Vizgen. The Takei et al. DNA seqFISH+ and sequential-immunofluorescence data are on Zenodo (doi:10.5281/zenodo.3735329) [33]. The PECAb data are on Zenodo (doi:10.5281/zenodo.8348965) [34].

## Code availability

Source code for the `spsheaf` package, the per-experiment scripts and the figure pipeline is publicly available at https://github.com/EduPH/spSheaf; the version used here, `spsheaf` 0.2.0, is archived on Zenodo (https://doi.org/10.5281/zenodo.24065558) and installable from PyPI (`pip install spsheaf`). The segmentation-free comparison (Section 2.3) is reproducible from the per-cell label tables and per-configuration metric tables tracked in that repository under `benchmarks/segfree/results/ (make matched-selection)`; the result tables behind every figure and table reported here, one row per configuration, are tracked under `reported/`, which lists for each file the claim it supports and the command that produced it, so any number above can be checked without re-running the pipeline (`make reported-check` verifies that the tracked copies still match the run outputs); the SpaceTx analyses are reproduced by `experiments/spacetx/` and `benchmarks/segfree/run spacetx.py (make spacetx spacetx-segfree spacetx-ficture)`; the raw competitor inputs are not redistributed.

## Acknowledgements

E.P-H. is supported by the European Union’s Horizon Europe research and innovation programme under the Marie Sklodowska-Curie grant agreement No. 101153039 (CHALKS), and by the IMUS–María de Maeztu grant CEX2024-001517-M, funded by MICIU/AEI/10.13039/501100011033. The authors are supported by JST CREST No. JPMJCR24Q1.

## Author contributions

E.P-H. designed the method, implemented the software, performed the experiments and benchmarks, analysed the results and wrote the manuscript. K.M. contributed to the interpretation of the results and the revision of the manuscript. Y.I. conceived the research problem, defined the study scope and datasets to be analysed, and contributed to the interpretation of the results and revision of the manuscript. All authors approved the final version.

## Extended Data

**Extended Data Table 1.**
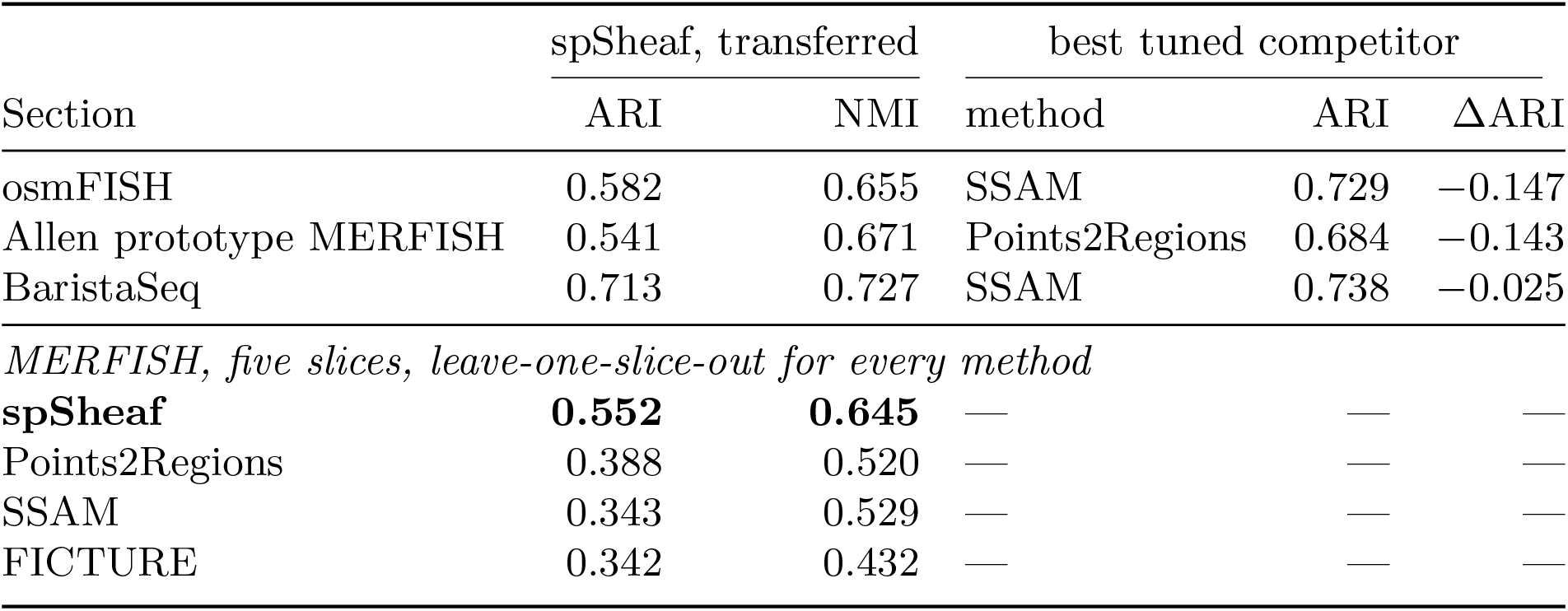
What the method does with no tuning. Table 1 tunes every method on the data it is scored on, which is the comparison the competitors’ published values demand but which measures none of them out of the box. This table asks the complementary question. *Top*: the three single sections, with every spSheaf parameter fixed on the MERFISH benchmark and carried over unchanged — the local coordinate reference and the *h, d, σ*_*s*_, *α*_*b*_, *α*_*s*_, *k*_graph_ and *k*_nbrs_ of Supplementary Table 2, with only the grid spacing and *k* set from the platform — against each section’s best competitor at the argmax of its own grid, selected on that section’s own annotation (the best competitor row of Table 1). Nothing about the section being scored enters the spSheaf configuration. Tuning, the reference included, adds 0.091 ARI on osmFISH and 0.213 and 0.044 on the two SpaceTx sections (Table 1). *Bottom*: the MERFISH benchmark under leave-one-slice-out, the strictest rule its five slices allow, where every method’s configuration is re-selected on the four slices it is not scored on and scored on the fifth (Fig. 2c, Section 4.12); spSheaf’s single development configuration applied to all five slices gives 0.571 and 0.655 instead. The identity of the leading competitor changes from section to section, so no single comparator is robust across the four.

| Section | spSheaf, transferred |  | best tuned competitor |  |  |
|---|---|---|---|---|---|
|  | ARI | NMI | method | ARI | ΔARI |
| osmFISH | 0.582 | 0.655 | SSAM | 0.729 | −0.147 |
| Allen prototype MERFISH | 0.541 | 0.671 | Points2Regions | 0.684 | −0.143 |
| BaristaSeq | 0.713 | 0.727 | SSAM | 0.738 | −0.025 |
| *MERFISH, five slices, leave-one-slice-out for every method* |  |  |  |  |  |
| **spSheaf** | **0.552** | **0.645** | — | — | — |
| Points2Regions | 0.388 | 0.520 | — | — | — |
| SSAM | 0.343 | 0.529 | — | — | — |
| FICTURE | 0.342 | 0.432 | — | — | — |

**Extended Data Fig. 1.**
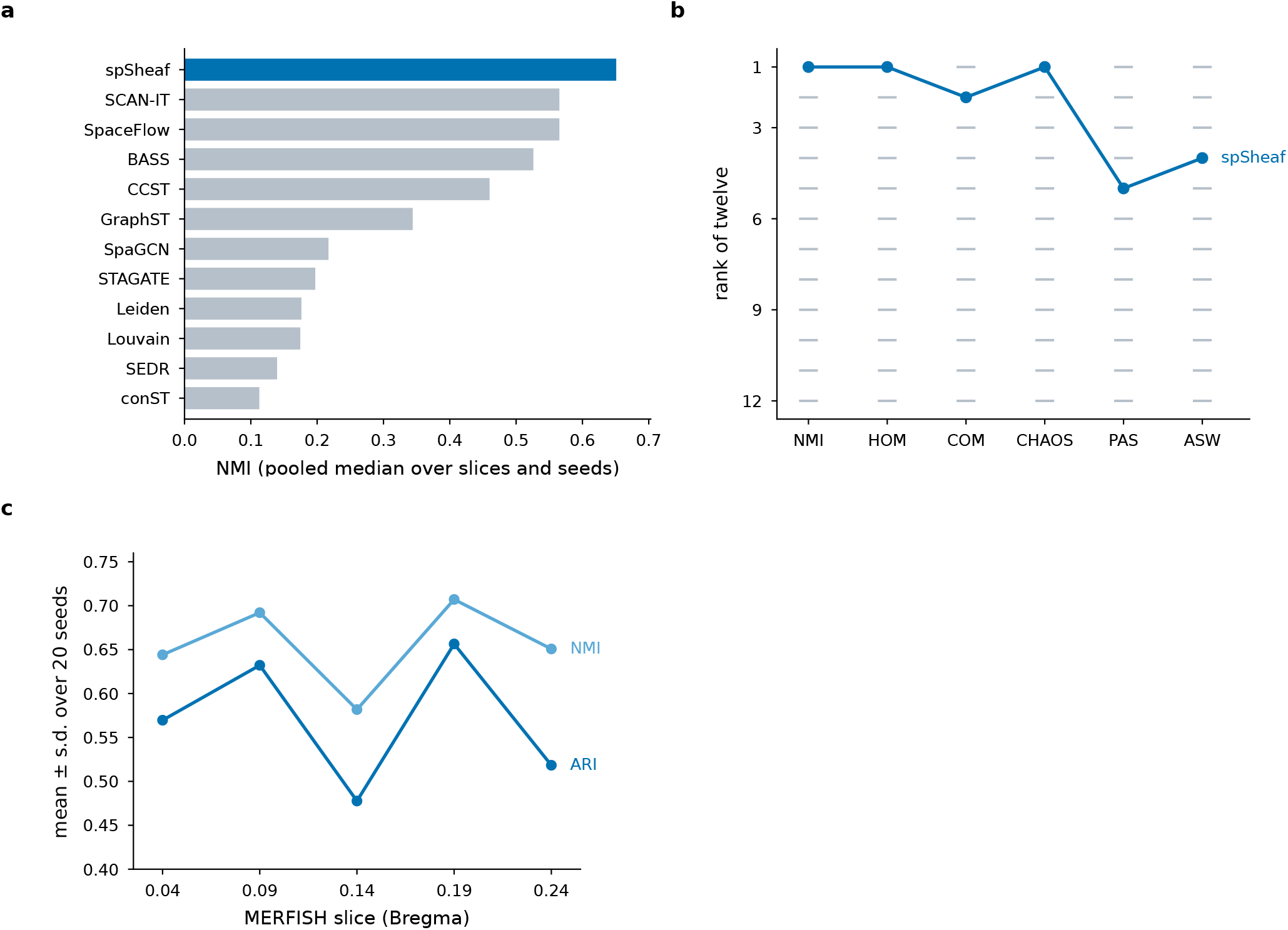
SDMBench ranking and seed sensitivity on the MERFISH benchmark. Rebuilt from the source data of Fig. 2d of ref. [24]. **a**, NMI for the twelve methods — the eleven baselines with MERFISH results, plus spSheaf — as pooled medians over slices and seeds. **b**, Rank of each method on the six ranked SDMBench metrics; horizontal marks are the eleven baselines and the line traces spSheaf. Moran’s I and Geary’s C are not ranked (Section 2.2). **c**, Mean *±* s.d. of ARI and NMI over 20 *k*-means seeds, per slice. Per-slice domain maps are shown in Supplementary Fig. 1.

**Extended Data Fig. 2.**
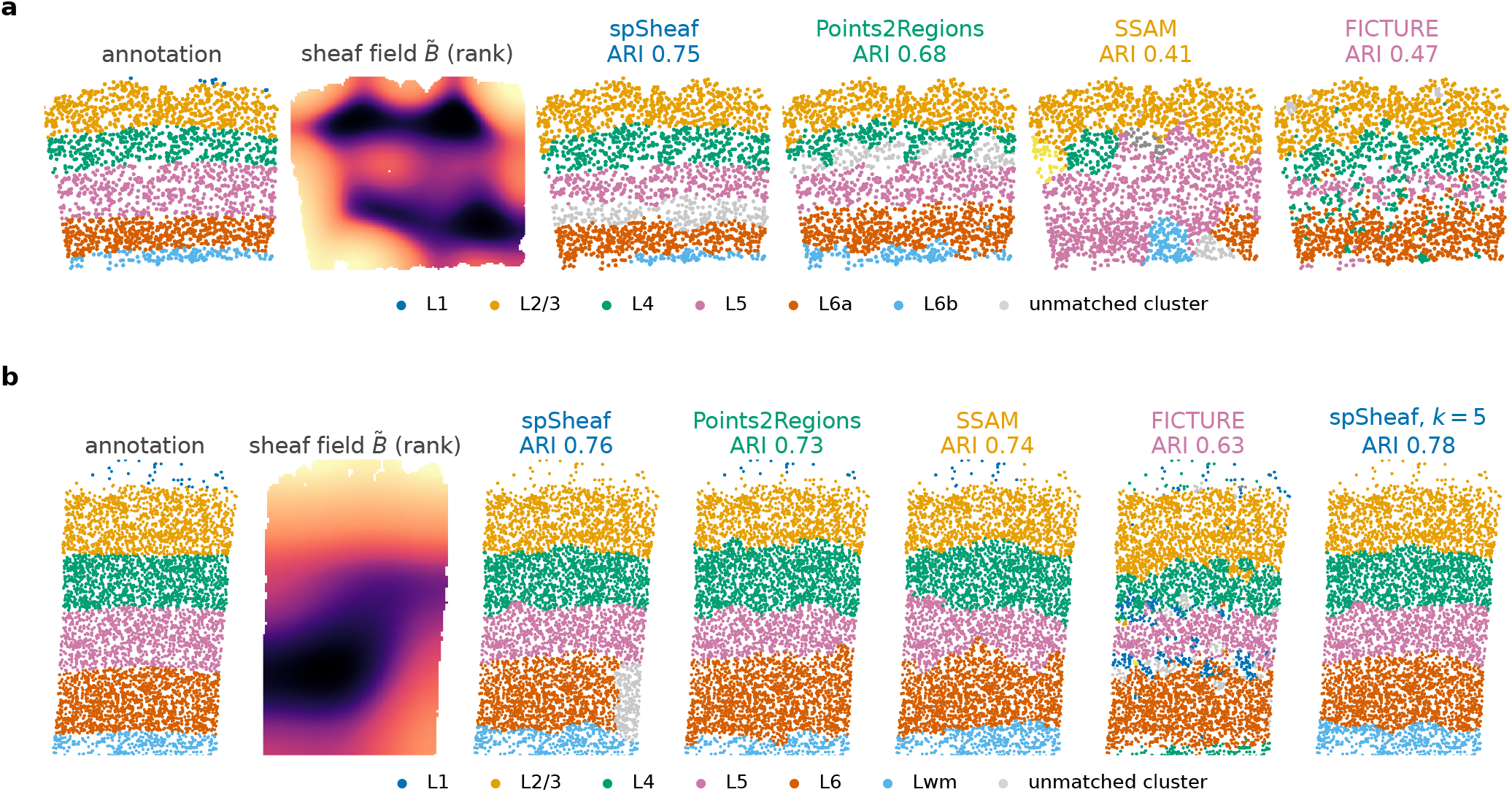
The two independent SpaceTx cortical sections, per annotated cell. **a**, Allen Institute prototype MERFISH (2,150 cells, six layers); **b**, BaristaSeq (4,432 cells, six classes). In each row: the layer annotation; the smoothed sheaf disagreement field 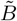 on the spSheaf grid, shown as its percentile rank among grid points under tissue (the tissue edge is orders of magnitude above the laminar transitions and would otherwise saturate the scale); and spSheaf, Points2Regions, SSAM and FICTURE, each at its own in-sample argmax on that section (Table 1). Labels are Hungarian-matched to the annotation; clusters beyond the annotated number (SSAM returns nine domains on **a**, FICTURE fifteen on **b**) are grey. The ARI above each map is the value of Table 1. On **a** spSheaf recovers L2/3 at recall 0.97 and L6b at 0.55; on **b** L5 and L6 at 0.80 and 0.79. Layer 1 (9 cells) is recovered by no method on **a**; on **b** (32 cells) Points2Regions, SSAM and FICTURE each recover about half of it (0.53, 0.50, 0.50) and spSheaf none. A cluster that shares no cell with the annotation class it is matched to is grey: on **a** every method has one such cluster, assigned to L1 by the matching; on **b** spSheaf’s sixth cluster is a *Synpr* -high patch within L6 (201 cells, identical over four seeds) and L1 is assigned to L2/3. The last column of **b** is the same configuration at *k* = 5, one below the annotated count: the patch merges into L6 and the five remaining layers are recovered at ARI 0.782.

**Extended Data Fig. 3.**
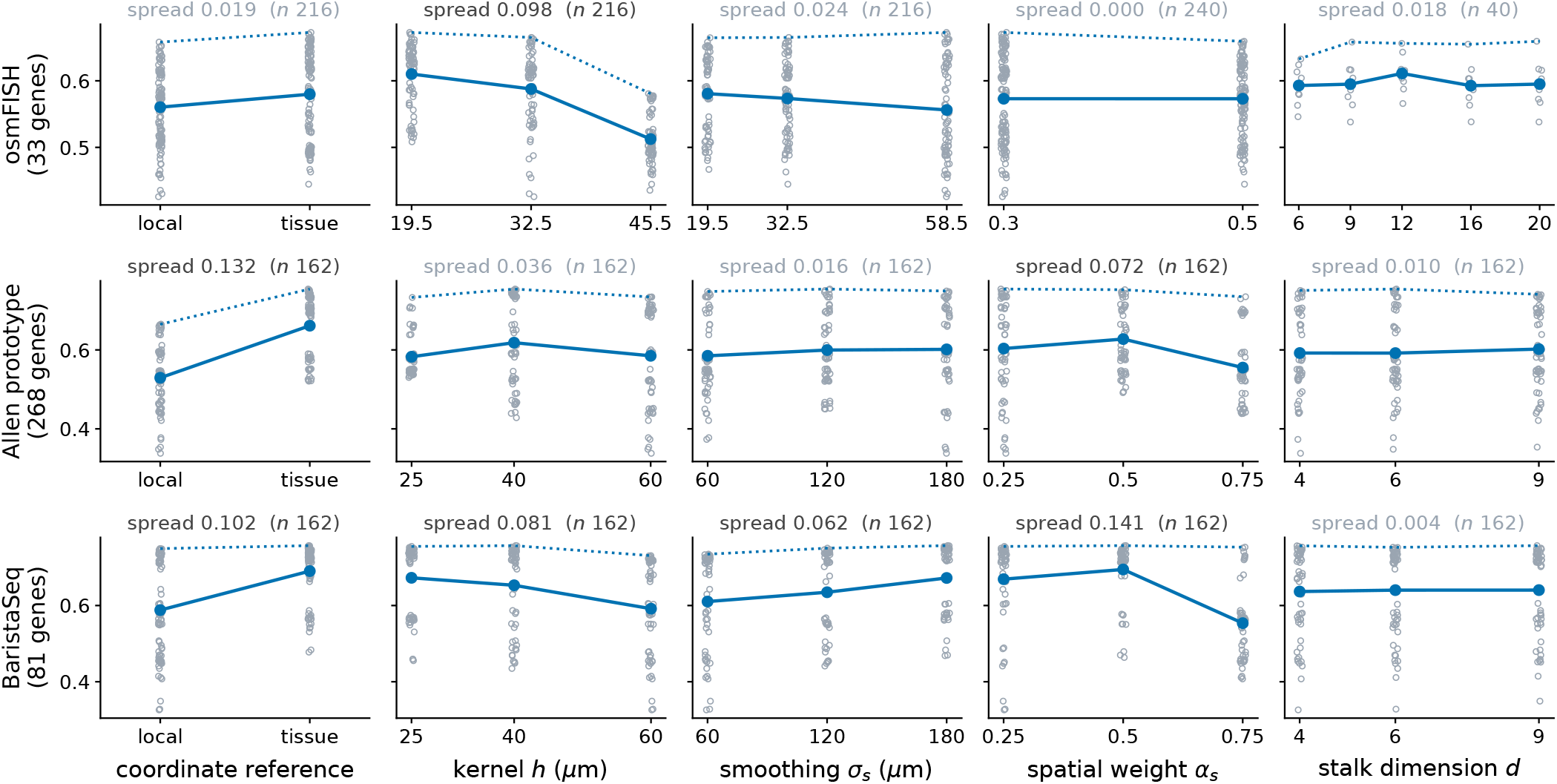
Which hyperparameters the accuracy depends on. ARI against each swept parameter on the three single sections, from the in-sample grids of Section 4.11 (osmFISH 240 configurations, the Allen prototype and BaristaSeq 162 each), marginalising over the remaining parameters: every configuration as a dot, the mean as the solid line and the maximum as the dotted line. Each marginal uses only the settings that were run at *every* level of the parameter shown, so that a parameter added on a sub-grid cannot inherit the others’ effect — unbalanced, the osmFISH stalk dimension appears to span 0.048 rather than 0.018, because its three highest levels were run only on a sub-grid. The number above each panel is the range of the mean, greyed where it is below 0.05, with the number of configurations entering it. No single parameter dominates everywhere. On the Allen prototype the largest effect is the coordinate reference, which moves the mean by 0.132, and on BaristaSeq it is second (0.102) to the spatial weight; the kernel bandwidth moves the mean by 0.036 and 0.081; on osmFISH the same reference is worth only 0.019 and the bandwidth leads at 0.098. The tissue reference is the better setting on all three, but by margins spanning an order of magnitude. The spatial weight ranges from decisive on BaristaSeq (0.141) to inert on osmFISH (0.000, the two levels indistinguishable once both references are pooled). Every one of these optima differs between sections, so both the reference and *σ*_*s*_ are selected per section (Section 2.6). The stalk dimension is the exception: over the swept range — 4–9 on the SpaceTx sections and 6–20 on osmFISH, the latter up to 57% of the 35-label feature space — the mean moves by 0.004–0.018 and the maximum by 0.004–0.027, with no monotone trend, so the local subspace is not short of capacity and the domains that are missed are not missed for want of dimensions (Section 2.5).

**Extended Data Fig. 4.**
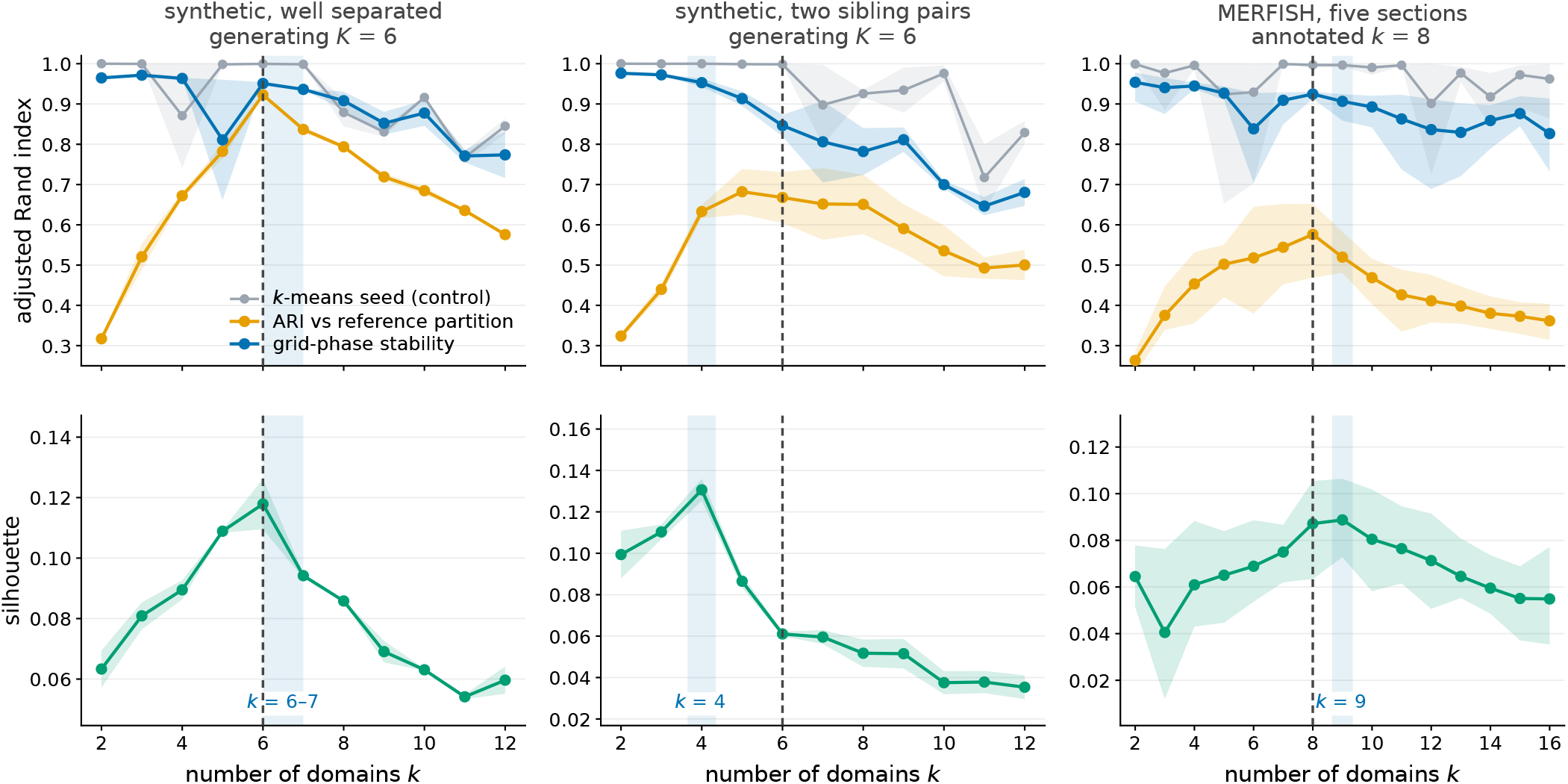
Annotation-free criteria for the number of domains *k*. Columns: a synthetic construction with *K* = 6 well-separated Voronoi domains, where *K* is known; a construction in which two pairs of domains are near siblings, where *K, K−*1 and *K −*2 are all defensible readings; and the five MERFISH sections, annotated with eight domains. Upper row, three quantities on the ARI scale: grid-phase stability (the ARI between a run at the default lattice phase and a run with the lattice offset by half a grid spacing, labels carried to the reference lattice by nearest neighbour), the *k*-means-seed agreement computed alongside as a control, and the ARI against the reference partition — the generating domains for the constructions, the annotation for MERFISH. Lower row, the silhouette of the normalised fibers over every grid point. Line, mean; band, the two independent draws (constructions) or the five sections (MERFISH). Dashed line, the reference *k*; shaded, the interval returned by the two criteria on the pooled curves — the silhouette argmax to the largest *k* whose stability is within 5% of the mean over *k ≤* 3. On the constructions the interval contains or under-counts *K* (6–7 and 6–8; 4–5 and 4–4). On the sections the stability bound is 16, 15, 8, 12 and 9 on the five slices, four of the ten lowest stability values coincide with a drop in the seed control, and the silhouette argmax is 10, 8, 9, 8 and 9; pooled, both criteria return 9 (Sections 2.7, 4.13).

**Extended Data Fig. 5.**
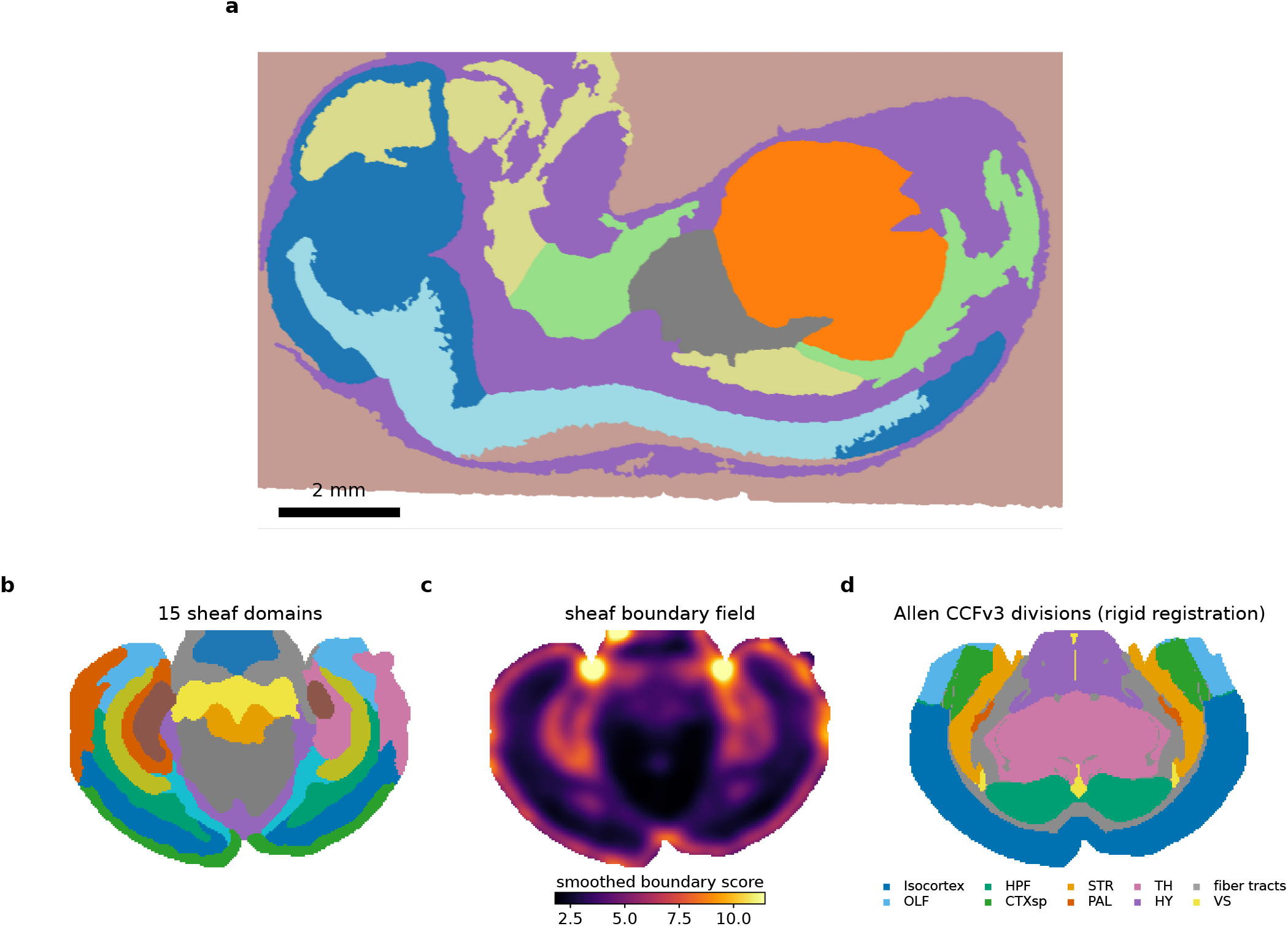
Scaling to whole-transcriptome, whole-section data. **a**, Sheaf domains over the whole Stereo-seq E16.5 mouse embryo (*∼*552 million transcripts) at 15 *µ*m grid resolution; eight organ-level compartments are resolved, with non-tissue captured as a single background domain. **b**, 15 sheaf domains on the Vizgen MERSCOPE mouse-brain coronal hemisection (54.7 million molecules; 50 *µ*m grid, 19,455 grid points), bilaterally symmetric and anatomically recognisable. **c**, The smoothed sheaf boundary field over the same grid. **d**, The ten major divisions of the Allen Common Coordinate Framework (CCFv3), rigidly registered onto the same grid, as an independent anatomical reference. Against these divisions over the 19,002 grid points inside the atlas (97.7% of the grid), the domains in **b** reach ARI 0.264 and NMI 0.460; after merging to matched divisions, accuracy 0.622 and ARI 0.425. The registration is rigid, so these values confound domain quality with registration error.

**Extended Data Fig. 6.**
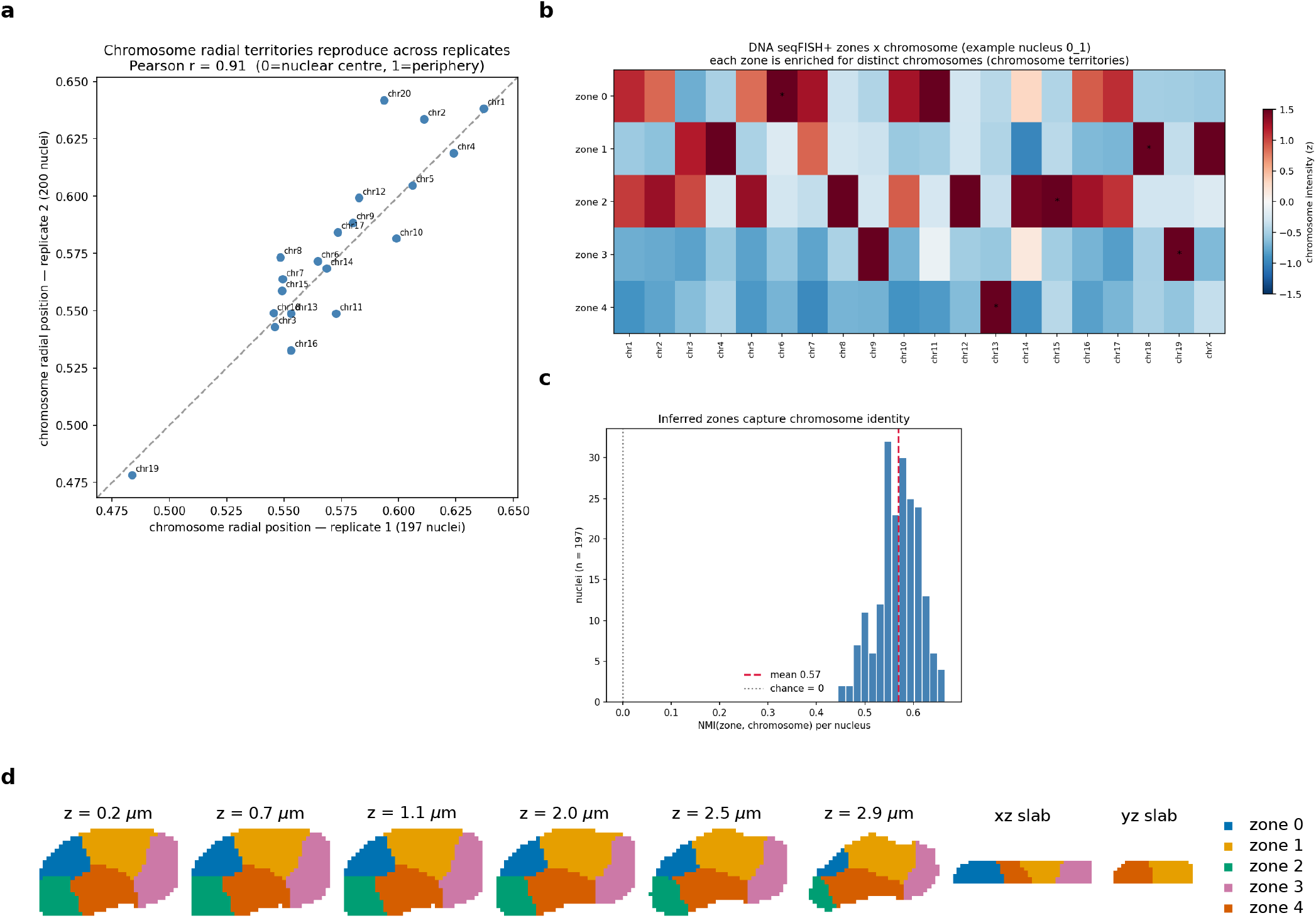
DNA seqFISH+ chromosome territories. **a**, Per-chromosome mean radial position (0 = nuclear centre, 1 = periphery) for two independent biological replicates (197 and 200 nuclei); the ordering reproduces across replicates at Pearson *r* = 0.91, with chromosome 19, the most gene-dense, the most interior in both replicates, consistent with the source study [33]; across the 19 chromosomes radial position and gene density are otherwise uncorrelated (*r* = *−*0.05). **b**, For an example nucleus, the zone *×* chromosome enrichment heatmap — each spatial zone is enriched for distinct chromosomes (star = dominant chromosome per zone), i.e. the zones recover chromosome territories. **c**, Distribution of the per-nucleus normalised mutual information between the inferred zones and chromosome identity across 197 nuclei (mean 0.57), confirming that the zones capture chromosome territories rather than arbitrary partitions. **d**, One nucleus, drawn from its voxel zone table: six optical sections at evenly spaced depths and two orthogonal slabs (xz at the median *y*, yz at the median *x*), coloured by the recovered three-dimensional chromosome-territory zone.

**Extended Data Fig. 7.**
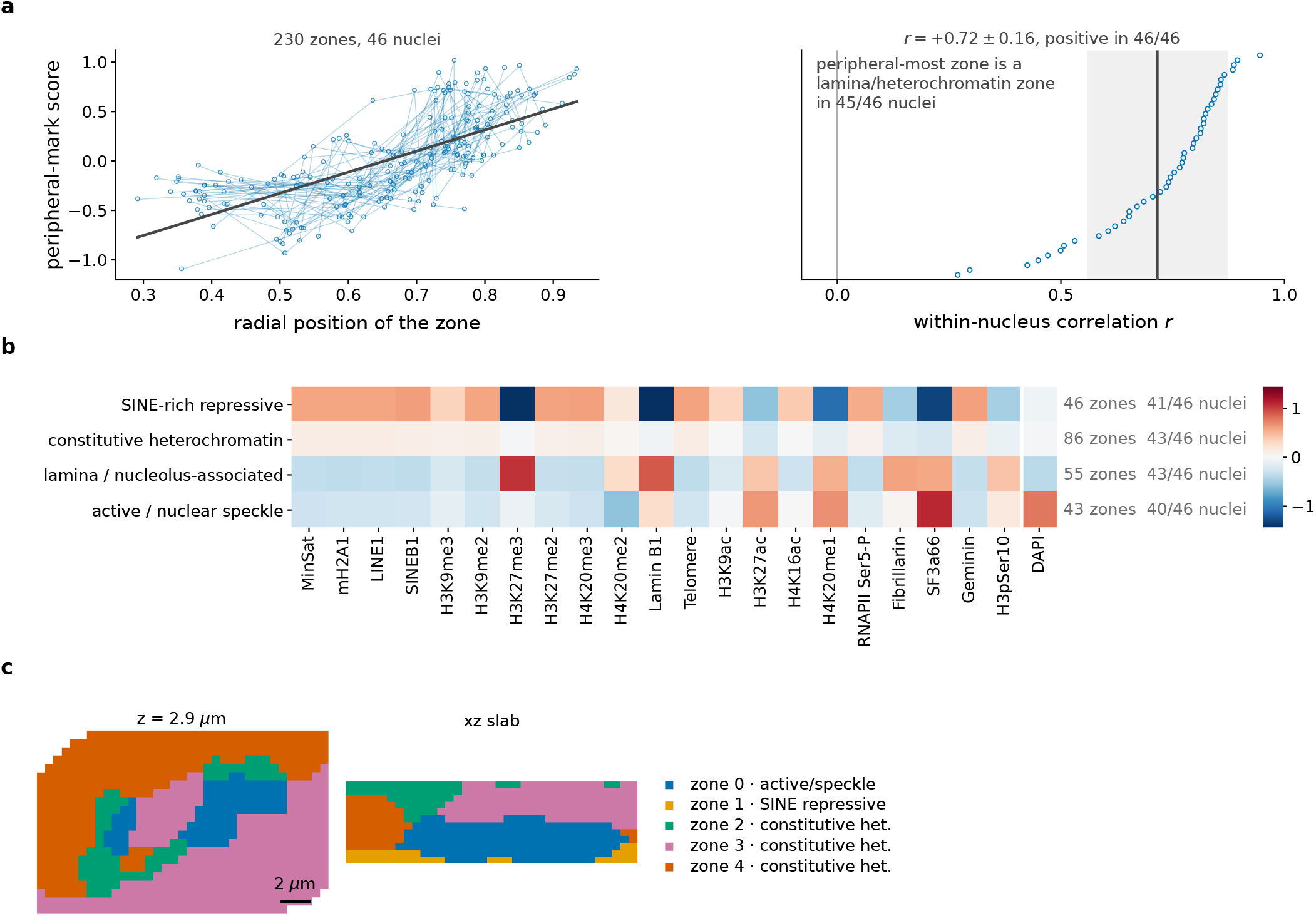
Chromatin-mark nuclear zones from sequential immunofluorescence. **a**, A zone’s peripheral-mark score against its radial position. Left: all 230 zones of all 46 nuclei, one line per nucleus, pooled fit in black. Right: the correlation computed within each nucleus, one point per nucleus, ordered; the black line and band are the mean and standard deviation. The within-nucleus correlation is *r* = +0.72*±*0.16 and positive in 46 of 46 nuclei — the claim holds inside every nucleus separately, which the pooled cloud alone could not establish — and the peripheral-most zone is a lamina/heterochromatin zone in 45 of 46, reproducing the known radial architecture of the nucleus. **b**, Chromatin-mark composition per zone archetype: the 230 zones pooled and clustered into recurring archetypes, profiles centred across marks before clustering (composition, not level). The number of archetypes is chosen on the data: the silhouette over *k* = 2–6 is 0.373, 0.275, 0.296, 0.283, 0.240, and *k* = 4 is the best choice above the trivial split (subsample stability 0.91). Each archetype recurs in 40–43 of the 46 nuclei and is named from the marks it is enriched for; columns are the 21 marks ordered heterochromatin *→* active *→* structural, DAPI at the right. The structure is a dominant heterochromatin-to-active axis with a secondary component: the constitutive-heterochromatin and SINE-rich profiles remain correlated at *r* = +0.79. **c**, One nucleus, drawn from its voxel zone table: a mid-depth optical section and one orthogonal slab, coloured by that nucleus’s own recovered zones (21-mark feature fiber), each labelled with the pooled archetype it falls in; three fall in the constitutive-heterochromatin archetype, so the zones are kept distinct rather than recoloured by archetype.

**Extended Data Fig. 8.**
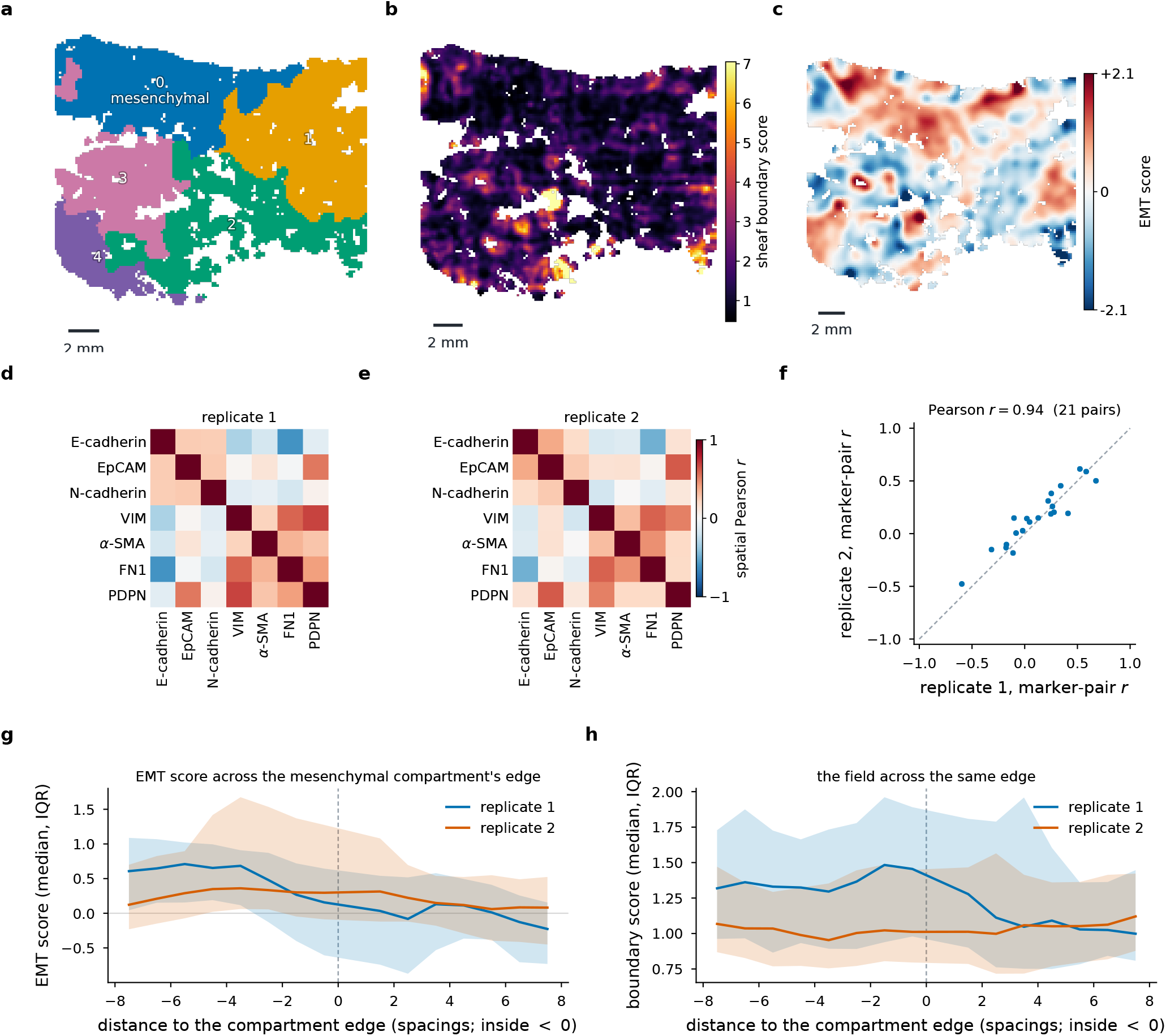
PECAb multiplexed protein imaging of a human uterine car-cinosarcoma. The feature-fiber variant applied to a non-transcriptomic modality (fiber = per-cell antibody intensities, 35-antibody panel, 7,657 cells). **a**, Five protein-state domains (within-domain edge fraction 0.968); domain 0 is a discrete mesenchymal/myofibroblast compartment (*α*-SMA, FN1, vimentin). **b**, The sheaf boundary field over the same section, delineating protein-state transitions. **c**, The continuous mesenchymal-minus-epithelial (EMT) score, a continuum of epithelial-to-mesenchymal states of which the compartment is the mesenchymal extreme (its edge is examined in **g**,**h**). **d**,**e**, Spatial co-localisation (Pearson *r*) of the lineage markers for two independent tissue sections: VIM, *α*-SMA and FN1 co-localise and are anti-correlated with E-cadherin in both replicates (*r* from *−*0.13 to *−*0.60); PDPN co-localises with VIM but also with EpCAM (0.52, 0.61), as expected for a marker present in both lineages here. **f**, Replicate 1 versus replicate 2 marker-pair correlations (21 pairs) agree at Pearson *r* = 0.94 (dashed line, identity), establishing that the recovered EMT structure is reproducible. Because the antibody panel is dominated by signalling-state (phospho) markers and the tissue is a fine epithelial–mesenchymal continuum, the output is a continuous axis plus one discrete compartment rather than sharply separated cell-type domains. No spatial-domain ground truth exists, so evaluation is qualitative and reproducibility-based. **g**,**h**, The EMT score (**g**) and the smoothed disagreement field (**h**) against the signed distance to the mesenchymal compartment’s edge, in grid spacings (150 px; negative inside), both replicates: median and interquartile range per spacing. In replicate 1 the score falls from +0.65 to *−*0.06 across the edge with a logistic width of one spacing, at the resolution of the 300-px kernel; in replicate 2 the compartment differs from its surroundings by 0.16 only. The field is 1.05*×* and 0.95*×* its interior median at the edge.

