## Supplementary information for "spSheaf: sheaf-theoretic spatial domain inference from raw molecules without cell segmentation"

**Supplementary Table 1** Per-slice performance of spSheaf on the Moffitt et al. (2018) MERFISH benchmark. Metrics: Adjusted Rand Index (ARI), Normalised Mutual Information (NMI), Homogeneity (HOM), Completeness (COM), CHAOS, Proportion of Arbitrarily Shaped clusters (PAS), Average Silhouette Width (ASW), and time. *Grid points* is the number of vertices of the spSheaf grid on that slice. Values are means over 20 random $k$-means seeds (the seed-sensitivity run of Extended Data Fig. 1c of the main text); the benchmark ranking (Extended Data Fig. 1a,b) uses pooled medians over all seeds and slices, so its per-metric values differ slightly.

| Slice | Grid points | ARI | NMI | HOM | COM | CHAOS | PAS | ASW | Time (s) |
|---|---|---|---|---|---|---|---|---|---|
| 0.04 | 14,255 | 0.570 | 0.644 | 0.657 | 0.632 | 0.0274 | 0.0294 | −0.018 | 29.0 |
| 0.09 | 14,181 | 0.632 | 0.692 | 0.702 | 0.683 | 0.0276 | 0.0284 | −0.048 | 29.3 |
| 0.14 | 14,327 | 0.478 | 0.582 | 0.591 | 0.573 | 0.0266 | 0.0257 | −0.062 | 30.2 |
| 0.19 | 14,293 | 0.656 | 0.707 | 0.714 | 0.700 | 0.0266 | 0.0247 | −0.023 | 29.4 |
| 0.24 | 14,376 | 0.518 | 0.651 | 0.668 | 0.634 | 0.0273 | 0.0317 | 0.007 | 29.6 |
| **Mean** | **14,286** | **0.571** | **0.655** | **0.666** | **0.644** | **0.0271** | **0.0280** | **−0.029** | **29.5** |

**Supplementary Table 2** MERFISH configuration.

| Symbol | Value | Description |
|---|---|---|
| $\delta$ | 15 $\mu$m | Grid spacing |
| $h$ | 25 $\mu$m | Gaussian kernel bandwidth |
| $k_{\text{graph}}$ | 6 | Spatial graph neighbours |
| $k_{\text{nbrs}}$ | 50 | Local PCA neighbourhood size |
| $d$ | 6 | Uniform stalk dimension |
| $\sigma_s$ | 60 $\mu$m | Boundary smoothing radius |
| $\alpha_b$ | 0.5 | Exponential weighting strength |
| $\alpha_s$ | 0.5 | Spatial coordinate weight |
| $k_{\text{clusters}}$ | 8 | Number of output domains |
| $\tau$ | 0.01 | Minimum fragment fraction |
| reference | local | Coordinate reference $\mu_v$, selected here; default: section mean $\bar{f}$ |

**Supplementary Table 3** Per-dataset settings and special preprocessing. Grid spacing $\delta$, bandwidth $h$ and boundary-smoothing radius $\sigma_s$ are in each dataset's native units. On the three single annotated sections the settings are the argmax of the grids of Methods (the coordinate reference is part of each grid; osmFISH's local-reference argmax is $h = 32.5\,\mu$m, $\sigma_s = 19.5\,\mu$m, $\alpha_s = 0.5$, ARI 0.658). $k_{\text{graph}}$ and $\alpha_b$ are held at the values of Supplementary Table 2 throughout, and $k_{\text{nbrs}}$ unless noted. The runs on the datasets without annotation use the tissue reference, except the IF zones.

| Dataset | $\delta$ | $h$ | $\sigma_s$ | $d$ | $\alpha_s$ | ref. | $k_{\text{clusters}}$ | Special preprocessing |
|---|---|---|---|---|---|---|---|---|
| MERFISH (Moffitt) | $15\,\mu$m | $25\,\mu$m | $60\,\mu$m | 6 | 0.5 | local | 8 | none |
| osmFISH | $9.8\,\mu$m | $19.5\,\mu$m | $58.5\,\mu$m | 9 | 0.3 | tissue | 11 | none |
| Allen MERFISH (SpaceTx) | $10\,\mu$m | $40\,\mu$m | $120\,\mu$m | 6 | 0.25 | tissue | 6 | none |
| BaristaSeq (SpaceTx) | $10\,\mu$m | $40\,\mu$m | $180\,\mu$m | 9 | 0.5 | tissue | 6 | none |
| Stereo-seq E16.5 (coarse) | $100\,\mu$m | $25\,\mu$m | $60\,\mu$m | 6 | 0.3 | tissue | 48 | top-300 genes, 10% subsample, $k_{\text{nbrs}} = 30$ |
| Stereo-seq E16.5 (fine) | $15\,\mu$m | $25\,\mu$m | $60\,\mu$m | 6 | 0.3 | tissue | 8 | top-300 genes, 50% subsample, $k_{\text{nbrs}} = 30$ |
| MERSCOPE brain | $50\,\mu$m | $100\,\mu$m | $200\,\mu$m | 6 | 0.5 | tissue | 15 | single $z$, top-300 genes, 25% subsample |
| DNA seqFISH+ (3D) | $0.55\,\mu$m | $0.8\,\mu$m | $0.9\,\mu$m | 6 | 0.5 | tissue | 5 | feature fiber (chromosome intensities), $k_{\text{nbrs}} = 40$ |
| IF chromatin zones (3D) | $0.55\,\mu$m | $0.8\,\mu$m | $0.9\,\mu$m | 6 | 0.5 | local | 5 | 21-mark feature fiber, per-voxel norm., $k_{\text{nbrs}} = 40$ |
| PECAb protein | 150 px | 300 px | 100 px | 6 | 0.5 | tissue | 5 | feature fiber (antibody panel), $k_{\text{nbrs}} = 40$ |

**Supplementary Table 4** Every division of an annotated domain at $k = 8$, 9 and 10 on the five MERFISH sections. A domain is divided when two clusters have it as their majority annotation; *share* is the fraction of the annotated domain in the larger and the smaller cluster, *purity* the fraction of each cluster inside the domain, *offset* the larger-minus-smaller centroid difference in $\mu$m. $r$ is the correlation over the 135 genes between this division's difference vector and that of the same domain on the section named in parentheses, the best of the non-hemispheric pairs; *hemispheric* marks a division into the two hemispheres ($|\Delta x| > 0.8$ mm), set aside. *Genes* are the four most enriched in the smaller half (Mann–Whitney, $q < 0.01$).

| Slice | $k$ | Domain | Share | Purity | Offset | $r$ (with) | Genes, smaller half |
|---|---|---|---|---|---|---|---|
| 0.04 | 8 | MPN | 0.60 / 0.30 | 0.86 / 0.53 | −148, +140 | +0.88 (0.09) | *Npas1 Coch Npy2r Slc18a2* |
| 0.04 | 9 | MPA | 0.48 / 0.35 | 0.95 / 0.95 | +169, +218 | +0.43 (0.24) | *Chat Aqp4 Lmod1 Myh11* |
| 0.04 | 9 | MPN | 0.61 / 0.30 | 0.85 / 0.51 | −147, +140 | +0.88 (0.09) | *Npas1 Coch Npy2r Slc18a2* |
| 0.04 | 10 | MPA | 0.44 / 0.27 | 0.95 / 0.88 | +297, +207 | +0.36 (0.24) | *Aqp4 Lmod1 Myh11 Calcr* |
| 0.04 | 10 | MPN | 0.61 / 0.30 | 0.85 / 0.55 | −151, +141 | +0.88 (0.09) | *Npas1 Coch Npy2r Slc18a2* |
| 0.09 | 8 | MPN | 0.68 / 0.19 | 0.77 / 0.52 | −45, +121 | +0.88 (0.04) | *Slc18a2 Npy2r Creb3l1 Esr1* |
| 0.09 | 9 | MPA | 0.52 / 0.36 | 0.83 / 0.88 | +1171, −117 | hemispheric | — |
| 0.09 | 9 | MPN | 0.66 / 0.19 | 0.85 / 0.52 | −44, +127 | +0.88 (0.04) | *Slc18a2 Npy2r Creb3l1 Esr1* |
| 0.09 | 10 | BST | 0.51 / 0.44 | 0.93 / 0.86 | −1314, −7 | hemispheric | — |
| 0.09 | 10 | MPA | 0.51 / 0.37 | 0.83 / 0.92 | +1187, −110 | hemispheric | — |
| 0.09 | 10 | MPN | 0.65 / 0.19 | 0.86 / 0.52 | −42, +122 | +0.88 (0.04) | *Slc18a2 Npy2r Creb3l1 Esr1* |
| 0.14 | 8 | MPA | 0.57 / 0.33 | 0.97 / 0.54 | +19, −434 | — | *Esr1 Calcr Coch Gda* |
| 0.14 | 8 | PVH | 0.47 / 0.09 | 0.37 / 0.44 | +418, −338 | — | *Nos1 Npy1r Pdgfra Myh11* |
| 0.14 | 9 | BST | 0.77 / 0.03 | 0.92 / 0.35 | +832, +108 | hemispheric | — |
| 0.14 | 9 | MPA | 0.56 / 0.34 | 0.98 / 0.56 | +12, −434 | +0.18 (0.24) | *Esr1 Calcr Gda Coch* |
| 0.14 | 10 | BST | 0.41 / 0.39 | 0.91 / 0.83 | +1280, −61 | hemispheric | — |
| 0.14 | 10 | MPA | 0.56 / 0.33 | 0.98 / 0.55 | +21, −426 | +0.24 (0.24) | *Esr1 Calcr Coch Gda* |
| 0.14 | 10 | PVH | 0.57 / 0.07 | 0.68 / 0.38 | +439, −158 | — | *Nos1 Creb3l1 Syt2 Npy1r* |
| 0.19 | 9 | BST | 0.52 / 0.40 | 0.79 / 0.90 | −132, +245 | +0.84 (0.24) | *Isl1 Esr1 Amigo2 Arhgap36* |
| 0.19 | 10 | BST | 0.52 / 0.39 | 0.82 / 0.90 | −162, +254 | +0.84 (0.24) | *Isl1 Esr1 Amigo2 Arhgap36* |
| 0.19 | 10 | MPA | 0.43 / 0.42 | 0.83 / 0.93 | −1272, −35 | hemispheric | — |
| 0.24 | 8 | BST | 0.54 / 0.33 | 0.91 / 0.90 | −90, +141 | — | *Isl1 Arhgap36 Amigo2 Fbxw13* |
| 0.24 | 9 | BST | 0.45 / 0.40 | 0.73 / 0.92 | −114, +185 | +0.84 (0.19) | *Isl1 Arhgap36 Amigo2 Gpr165* |
| 0.24 | 9 | MPA | 0.87 / 0.06 | 0.78 / 0.58 | −457, −253 | +0.43 (0.04) | *Myh11 Lmod1 Fn1 Ccnd2* |
| 0.24 | 10 | BST | 0.45 / 0.40 | 0.71 / 0.90 | −131, +192 | +0.84 (0.19) | *Isl1 Arhgap36 Gpr165 Amigo2* |
| 0.24 | 10 | MPA | 0.58 / 0.05 | 0.89 / 0.56 | −557, −357 | +0.36 (0.04) | *Myh11 Lmod1 Fn1 Ccnd2* |

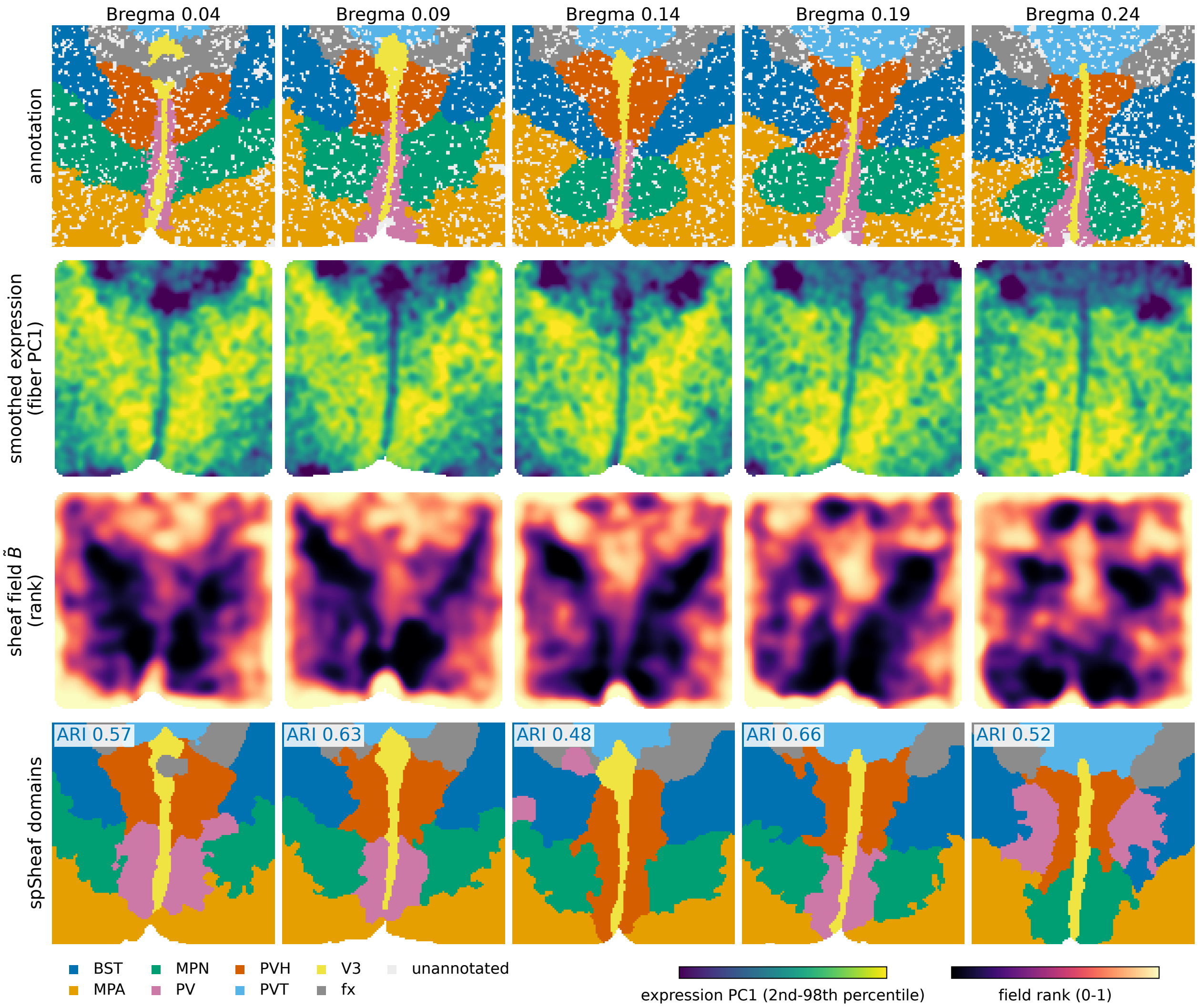


**Supplementary Fig. 1 Per-slice fields and domains on the MERFISH benchmark.** One row per slice (Bregma 0.04–0.24), every panel drawn from the same cache the manuscript's numbers are computed on. Columns: the annotation, at grid level, with unannotated grid points in grey; the smoothed expression the sheaf is built on, as the first principal component of the fibers; the smoothed disagreement field $\tilde{B}$, as a percentile rank; and the recovered domains, Hungarian-matched to the annotation. The ARI on each row is that slice's value in Supplementary Table 1, the 20-seed cell-level mean; the map drawn is one seed scored at grid level, which differs by up to 0.02. Both continuous fields are shown on the interior of the section only: at its edge the fiber neighbourhood is one-sided and the disagreement is far above any internal transition, which would otherwise set the colour scale. The field is elevated over the transitions between programs rather than sharply at the annotated interfaces — Section 2.4 of the main text quantifies that width, and it is the extent the exponential weight acts on.

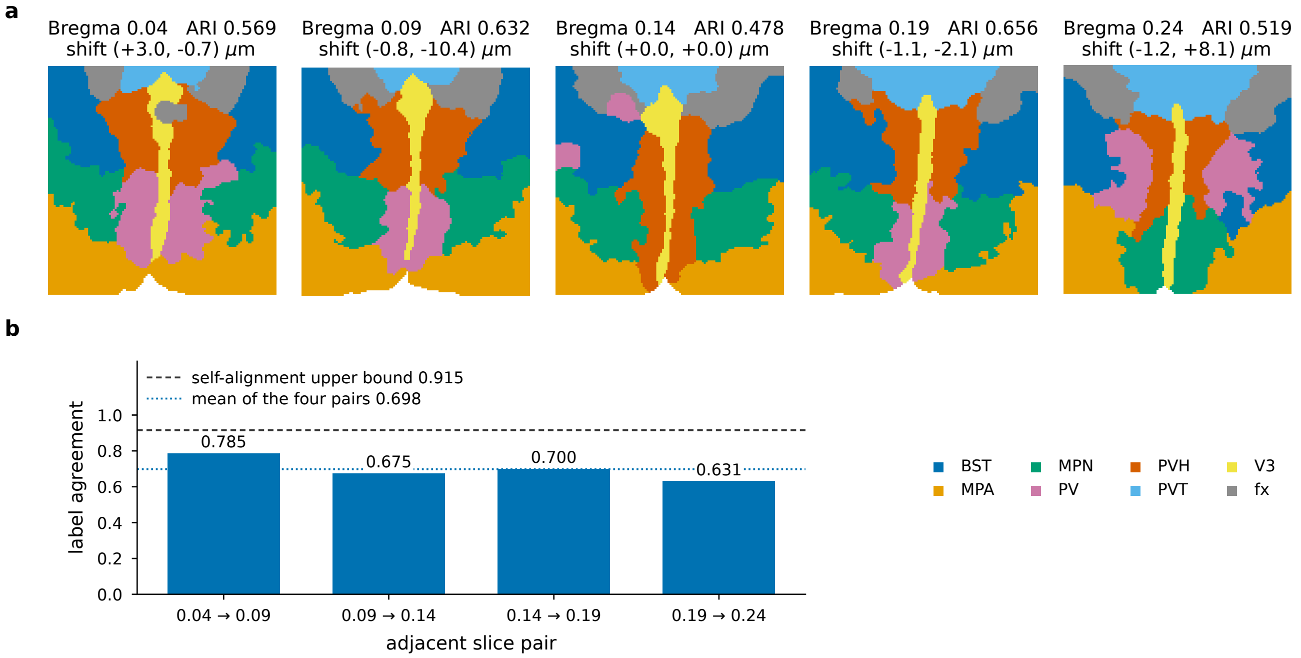


**Supplementary Fig. 2 Multi-slice alignment across serial sections.** **a**, The five consecutive MERFISH slices after gene-centroid registration, drawn in the common frame and coloured by domain (labels Hungarian-matched to the annotation; per-slice ARI and the recovered translation above each). **b**, Cross-slice label agreement for the four adjacent pairs, computed over inter-slice edges within 45 $\mu$m after registration and Hungarian matching of the two slices' labels; the dashed line is the self-alignment upper bound, a slice registered against a shifted copy of itself. Agreement is well above chance on every pair, confirming that per-slice predictions are spatially coherent across depth.

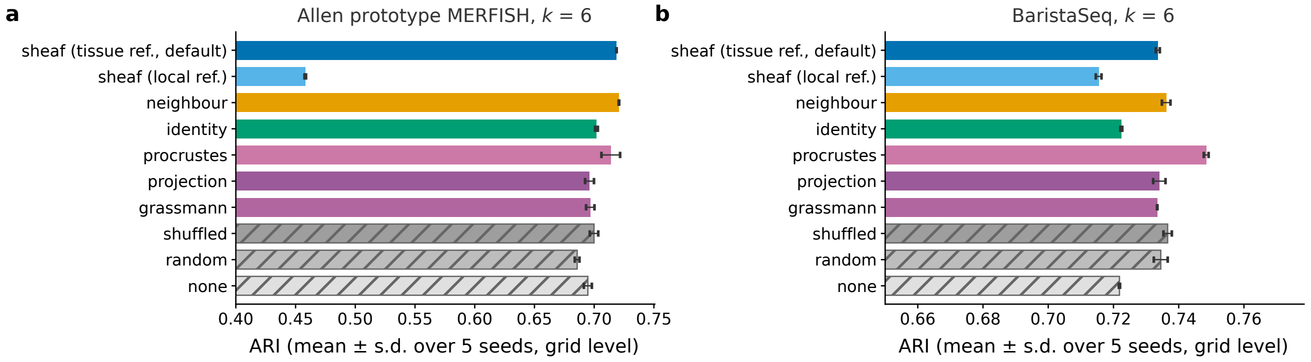


**Supplementary Fig. 3 The edge-score ablation on the two SpaceTx sections.** The ten arms of Fig. 2d of the main text, run on the Allen prototype MERFISH (**a**) and BaristaSeq (**b**) sections at each section's selected configuration (tissue coordinate reference; $h = 40\,\mu$m; $\sigma_s = 120$ and $180\,\mu$m; $d = 6$ and 9; $\alpha_s = 0.25$ and 0.5; $k = 6$), scored at grid level against the annotation carried to the grid by the nearest registered cell. Bars are means over five $k$-means seeds, whiskers the standard deviation; hatched bars are the three null controls. On the Allen prototype the sheaf score reaches 0.719 against 0.721 for the kernel baseline and 0.695 for no weighting, and the local-reference score 0.458; on BaristaSeq 0.734 against 0.736, 0.722 and 0.715, with the Procrustes residual at 0.748. Neither section separates the alignment-adjusted score from the kernel baseline.
